# Biodiversity knowledge and conservation shortfalls in Rivulidae fishes

**DOI:** 10.64898/2026.07.04.736495

**Authors:** João Henrique Alliprandini da Costa, Gustavo Henrique Soares Guedes

## Abstract

The biodiversity crisis is exacerbated by persistent gaps in taxonomic, geographic, and conservation knowledge. This study provides a comprehensive assessment of Linnean, Wallacean, and conservation shortfalls in Rivulidae (Cyprinodontiformes), one of the most diverse families of Neotropical freshwater fishes. To this end, an extensive dataset was compiled, comprising 494 valid species, 51 synonyms, 3,419 occurrence records, and information on life cycle, distribution, and conservation status. The Linnean shortfall remains open: after 1975, the rate of species description increased sharply, and estimates indicate that 103 species remain undescribed (95% CI: 53–212). The year of species description was influenced by detectability and accessibility factors, with larger species, more widely distributed species, and species occurring in more densely populated areas being described earlier. The Wallacean shortfall was broad and spatially uneven: only 5.34% of the region was considered adequately sampled. The conservation shortfall was also substantial: 179 species are threatened with extinction, 69 species remain Not Evaluated, and 62 are Data Deficient. The mean time between taxonomic description and first IUCN extinction-risk assessment was 25.7 years. Moreover, 59.9% of species have no records within protected areas, including 69.3% of threatened species. These findings synthesize an urgent challenge: biodiversity knowledge and conservation shortfalls must be overcome simultaneously to protect species that are still being discovered, remain poorly documented spatially, and are restricted to habitats under intense anthropogenic pressure. The conservation of Rivulidae cannot wait for complete knowledge; action amid uncertainty is necessary to prevent both known and unknown species from disappearing.

## 1. Introduction

Our knowledge of Earth’s biodiversity remains profoundly incomplete. Despite centuries of research, biodiversity information is unevenly distributed across taxa, regions, and ecosystems, and these gaps are unlikely to be fully resolved in the foreseeable future (Hortal et al. 2015). Although only a small fraction of the world’s estimated species diversity has been formally described (Mora et al. 2011; Costello et al. 2013), our ignorance extends far beyond species identity. Major gaps also persist in our understanding of species distributions, evolutionary history, population abundance, functional traits, ecological interactions, and environmental tolerances (Hortal et al. 2015). Importantly, the information that is currently available is itself spatially and taxonomically biased, limiting our ability to accurately describe biodiversity patterns, prioritize conservation actions, and predict biological responses to ongoing environmental change (Mace 2004; Hortal et al. 2015; Frateles et al. 2026). This creates a fundamental conservation challenge. Effective conservation decisions cannot wait until biodiversity inventories become complete, particularly because many species may disappear before they are even discovered or formally described (Tedesco et al. 2014). Consequently, identifying and quantifying the dimensions of our ignorance has become as important as documenting biodiversity itself. In the era of biodiversity big data, comprehensive syntheses that explicitly measure biodiversity knowledge shortfalls while integrating available species information provide an essential framework for identifying research priorities and supporting more effective, evidence-based conservation planning (Hortal et al. 2015; Guedes et al. 2026a).

Among these biodiversity knowledge shortfalls, the Linnean shortfall refers to the proportion of extant and extinct species that remain undescribed (Hortal et al. 2015) and addresses one of the most fundamental questions in biodiversity science: how many species inhabit our planet (Lima et al. 2026). Reducing the Linnean shortfall is particularly important because progress toward virtually every other biodiversity knowledge dimension depends on species identity (Diniz-Filho et al. 2023; Guedes et al. 2026a). Another major dimension is the Wallacean shortfall, defined as the lack of information on species geographic distributions (Hortal et al. 2015). Because occurrence records represent one of the most accessible and widely available sources of biodiversity information (Frateles et al. 2026), understanding and reducing this shortfall is fundamental for accurately estimating species’ geographic ranges, describing biodiversity patterns, predicting future distributions, and supporting extinction-risk assessments (Hortal et al. 2015; Frateles et al. 2026).

Beyond biodiversity knowledge shortfalls, there is also growing recognition of a conservation shortfall, representing the gap between acquiring biodiversity knowledge and translating that knowledge into effective conservation actions. Although this dimension was not formally included among the biodiversity shortfalls proposed by Hortal et al. (2015), it has become increasingly relevant because many recently described species remain unevaluated for extinction risk or occur outside protected areas long after their scientific recognition (Rodrigues et al. 2004; Tapley et al. 2018). Quantifying these delays is crucial because conservation assessments and protected-area coverage underpin conservation prioritization, funding allocation, and policy implementation (Brooks et al. 2004). Consequently, measuring the time elapsed between species description and conservation assessment, together with the representation of species occurrences within protected areas, provides an objective framework for identifying species and regions where conservation actions have failed to keep pace with advances in taxonomic knowledge (Tapley et al. 2018).

Importantly, these knowledge and conservation shortfalls are not random. Patterns of scientific knowledge are strongly influenced by taxonomic and public interest, with charismatic or less threatened organisms generally receiving disproportionate research attention (Mammola et al. 2023; Mouquet et al. 2024). Likewise, intrinsic species characteristics, such as habitat use, geographic range size, and body size, can influence research effort (Mammola et al. 2023). Extrinsic factors also play an important role, including the difficulty of sampling particular environments, such as high-elevation regions (Reis et al. 2024; Frateles et al. 2026), as well as landscape characteristics affecting accessibility, including road density, river accessibility, proximity to biodiversity institutions, protected areas, human population density, and socioeconomic development (Moura et al. 2018; Penhacek et al. 2025). Therefore, beyond quantifying biodiversity knowledge and conservation shortfalls, it is equally important to understand the factors driving these biases, allowing future sampling and conservation efforts to be directed to species that need the most.

The family Rivulidae (Cyprinodontiformes) provides an exceptional model for assessing these shortfalls. First, Rivulidae is the seventh most species-rich family of freshwater fishes worldwide, comprising 494 valid species (Fricke et al. 2026a), and combines high species richness with an active taxonomic history (e.g., Alonso et al. 2023; Volcan et al. 2025; Abrantes et al. 2026). This makes the family particularly suitable for evaluating how taxonomic knowledge has accumulated over time, which uncertainties still persist, and how much diversity may remain undescribed. Second, unlike the most species-rich freshwater fish families in the Neotropical region (e.g., Loricariidae, Acestrorhamphidae), rivulids, as a rule, do not occupy large water bodies, such as major rivers and lakes, but instead exhibit high levels of endemism associated with spatially small, fragmented, shallow, or temporary aquatic habitats (Furness 2016; Loureiro et al. 2018; Costa et al. 2025), environments that have historically been neglected in ichthyological surveys. Third, a substantial proportion of Rivulidae (∼260 spp.) has an annual life cycle (Guedes et al. 2025a), characterized by the deposition of eggs in the substrate and the occurrence of embryonic diapause, a drastic, reversible, and regulated interruption of development associated with metabolic depression and molecular reorganization (Arezo et al. 2017; Zajic and Podrabsky 2020). Annual species develop and reproduce in temporary habitats that dry out during the dry season, eliminating the entire adult population (Furness 2016; Clouser et al. 2025). Diapausing embryos persist cryptically in egg banks buried in the sediment for months without access to liquid water, until hatching during the following rainy season (Berois et al. 2012). This life cycle adds a temporal dimension to the Wallacean shortfall, because the absence of records may reflect surveys conducted outside the hydrological window in which adults are detectable. Fourth, Rivulidae is subject to an unparalleled extinction threat (Volcan and Lanés 2018; Castro and Polaz 2020; Guedes and Araújo 2026). In Brazil, Rivulidae alone includes more threatened species than all other continental fish families combined: 131 versus 129 species across 23 families (MMA No. 1,667 2026). Together, these characteristics make Rivulidae an exceptional system for examining the uneven accumulation of taxonomic, geographic, and conservation knowledge across time and space.

Therefore, the main objective of this study is to provide an integrated assessment of the major taxonomic, geographic, and conservation knowledge shortfalls in Rivulidae. To this end, the most comprehensive dataset currently available for the family was compiled, integrating taxonomic information, occurrence records, geographic and biological attributes, and species threat status. The study is guided by three main sets of questions: (1) Linnean shortfall: how have the discovery of valid species and the production of synonyms varied over time? What is the estimated total richness of the family, considering both described species and those potentially awaiting description? Which geographic, environmental, biological, and anthropogenic factors help explain variation in the year of species description? Does the year of description vary spatially across the Neotropical region? (2) Wallacean shortfall: how is geographic knowledge of Rivulidae distributed? Where are well-sampled areas, spatial inventory gaps, and occurrence areas isolated from the main sampling nuclei? (3) Conservation shortfall: what is the current conservation status of Rivulidae species? How long is the time lag between taxonomic description and the first formal extinction risk assessment? To what extent are species, particularly threatened species, represented within protected areas, and is this representation concentrated mainly in strictly protected areas or in multiple-use areas? By integrating these three dimensions, this study aims to identify where knowledge of Rivulidae remains incomplete and to translate these shortfalls into applied evidence to guide future field surveys, taxonomic effort, extinction risk assessments, and conservation policies aimed at protecting Rivulidae diversity.

## 2. Methods

### 2.1. Taxonomic information

Taxonomic information for Rivulidae, including valid species, synonyms, original description dates, and authorship of species descriptions, was obtained from Eschmeyer’s Catalog of Fishes (Fricke et al. 2026a; accessed June 2026). The only exception was *Kryptolebias caudomarginatus* Seegers, 1984, treated as valid in Eschmeyer’s Catalog of Fishes but synonymized with *K. ocellatus* (Hensel, 1868) by Costa (2011). Because this synonymy has been adopted by Brazilian Rivulidae specialists (Pavanelli et al. 2024a), *K. caudomarginatus* was treated as a synonym of *K. ocellatus* in the dataset. The final dataset comprised 545 species names, including 494 currently recognized valid species and 51 synonyms.

### 2.2. Geographic distribution

To determine the known geographic distribution of each of the 494 valid species, occurrence records were compiled from multiple sources. First, all available records for 261 annual Rivulidae species were assembled from Guedes et al. (2025a). This dataset was complemented with records of non-annual Brazilian Rivulidae obtained from the Sistema de Avaliação do Risco de Extinção da Biodiversidade (SALVE) (ICMBio 2026a), an expert-validated platform containing information on species distributions. For species occurring outside Brazil and not represented in Guedes et al. (2025a) or SALVE–ICMBio, occurrence records were obtained from original species descriptions (e.g., Volcan et al. 2024; Alonso et al. 2025) and the Global Biodiversity Information Facility (GBIF 2026), retaining only records classified as “Material Citation”, “Material Sample”, or “Preserved Specimen” to minimize potential taxonomic errors associated with human observations. Records assigned to synonymous names were reassigned to their corresponding valid species. Duplicate occurrences were subsequently removed with the *CoordinateCleaner* R package (Zizka et al. 2019), and manually verified the geographic consistency of each record against the known country-level distribution of each species. The final dataset, representing the most comprehensive occurrence compilation assembled to date for the family, comprised 3,419 occurrence records (Fig. 1), covering 494 valid species and 49 genera of Rivulidae.

**Fig. 1.**
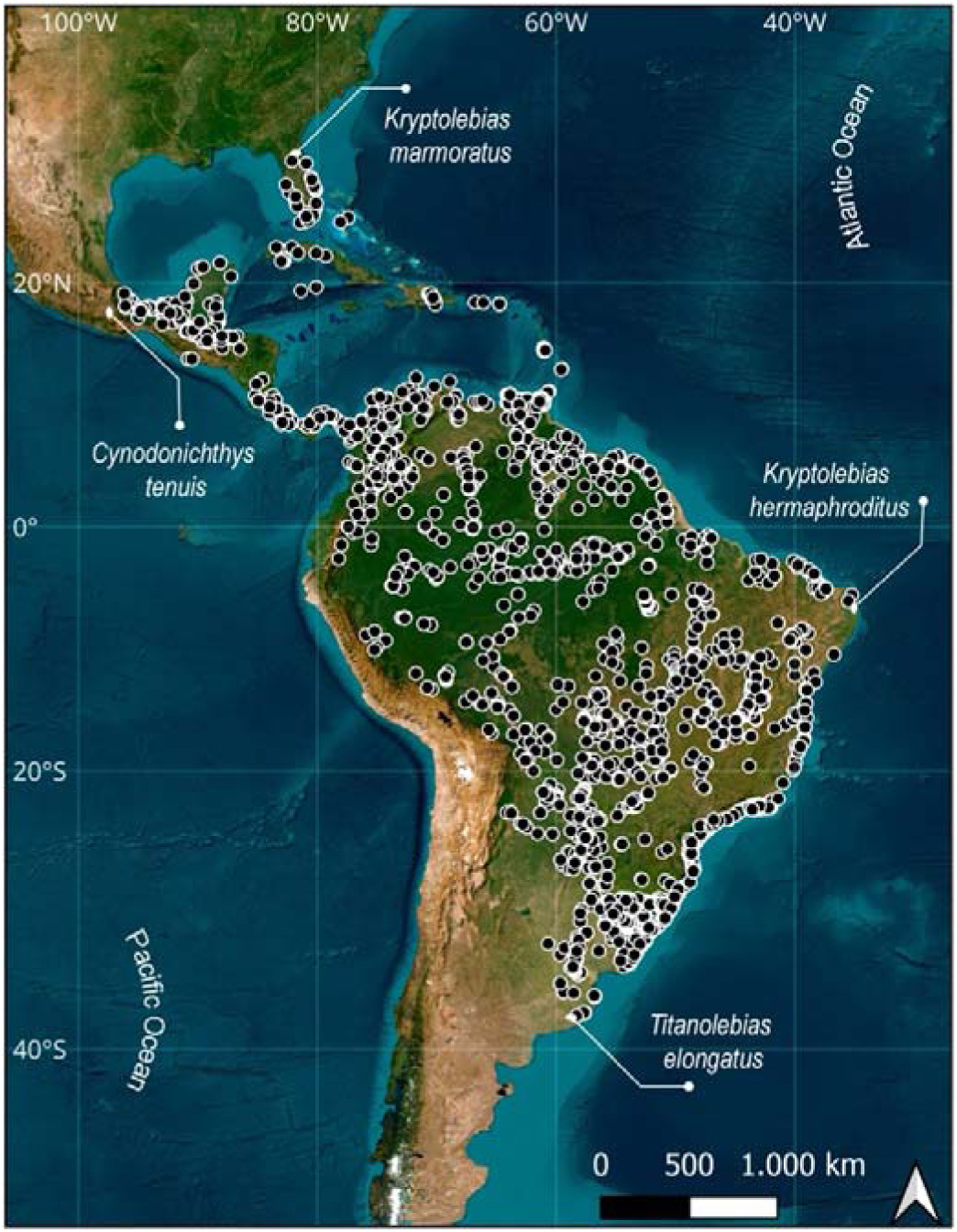
Spatial distribution of Rivulidae species (Cyprinodontiformes) in the Americas. The map indicates the geographic limits of the family’s distribution: north, *Kryptolebias marmoratus* (Poey 1880) in Florida, USA (29°16′02.0″N, 82°08′01.6″W); south, *Titanolebias elongatus* (Steindachner 1881) in Buenos Aires, Argentina (37°50′34.1″S, 58°17′17.9″W); east, *Kryptolebias hermaphroditus* Costa 2011 in Rio Grande do Norte, Brazil (5°40′25.9″S, 35°14′14.5″W); and west, *Cynodonichthys tenuis* Meek 1904 in Oaxaca, Mexico (18°09′58.1″N, 96°05′52.1″W).

### 2.3. Conservation status

The conservation status of species was compiled from the IUCN Red List of Threatened Species (IUCN, 2026). In addition, these data were supplemented and updated using assessments available through the Brazilian SALVE system (ICMBio 2026a), because: (i) Brazil follows the same categories and criteria adopted by the IUCN Red List (Souza et al. 2008); (ii) the conservation status of aquatic species occurring in the country was recently updated (MMA No. 1,667 2026); and (iii) regional assessments of Brazilian endemic species are frequently used by the IUCN to inform updates of the global Red List. When discrepancies existed between the global IUCN assessment and the Brazilian national assessment, the category based on the most recent assessment and/or the assessment most appropriate to the species’ geographic distribution was adopted (i.e., for species occurring in more than one country, the global IUCN category was retained). Species were assigned to four main conservation-status classes following Borgelt et al. (2022): (i) threatened species, including those classified as Critically Endangered (CR), Endangered (EN), or Vulnerable (VU); (ii) non-threatened species, including those classified as Near Threatened (NT) or Least Concern (LC); (iii) Data Deficient species (DD); and (iv) Not Evaluated species (NE). The DD and NE categories were treated separately because they represent distinct types of knowledge shortfalls. Data Deficient species have undergone formal assessment but lack sufficient information to reliably estimate their extinction risk, whereas Not Evaluated species have not yet been formally assessed (Borgelt et al. 2022; Cazalis et al. 2023).

### 2.4. Linnean Shortfall

To evaluate temporal patterns associated with the Linnean shortfall, the accumulation of valid species and synonyms through time was first quantified. The cumulative number of valid species and synonymous names by year of description was calculated, and the temporal change in the proportion of synonyms relative to valid species was also measured. The overall mean annual description rate across the full study period was also calculated, as well as description rates per decade for valid species. These analyses were used to describe temporal trends in species discovery and taxonomic synonymy before fitting predictive models of total species discovery. To estimate the total number of Rivulidae species, including both described and potentially undescribed species, the species discovery modelling framework used in recent analyses of biodiversity shortfalls was followed (Guedes et al., 2026a). Only valid species were used, and description data were grouped into 5-year intervals. For each interval, the number of species described during the interval (ΔSL), the cumulative number of species described before the interval (SL), the mean year of description, and the number of taxonomists involved in species descriptions during that interval (TL) were calculated. Author names were standardized, split into individual taxonomists, and counted as unique authors within each 5-year interval. Three species discovery models were fitted: the Lu and He model (Lu and He 2017), in which description efficiency varies as a function of both taxonomic effort and the number of species described per interval; the Joppa et al. model (Joppa et al. 2011), in which description efficiency varies as a function of taxonomic effort and time; and a logistic model, in which description efficiency varies as a function of the cumulative number of described species. Models were fitted using the gnls function from the *nlme* R package (Pinheiro et al. 2020), with residual variance modelled as a power function of the fitted values when convergence allowed, to account for over-or underdispersion (Guedes et al., 2026a). Models were ranked using the Akaike Information Criterion (AICc), and model support was assessed using AICc weights (wAICc). Uncertainty around total richness estimates was assessed using a bootstrap procedure with 100 resampled datasets. Following Guedes et al. (2026a), a negative exponential model was also evaluated. However, this model consistently failed to provide reliable parameter estimates, returning the starting value of total richness across all bootstrap replicates, and was therefore excluded from model comparisons.

Species-level information associated with the year of species description was then evaluated. For this analysis, a Bayesian Gamma regression model with a log link was fitted using the *brms* R package (Bürkner 2017). The response variable was the year of species description. Predictor variables included biological, geographic, environmental/hydrographic, and anthropogenic-accessibility predictors potentially related to species detectability, accessibility, collection probability, and therefore to variation in the year of species description (Table 1). Range size, body size, population density, road density, river order, and elevation were log10(x + 1)-transformed and subsequently standardized to mean 0 and standard deviation 1, to allow comparison of estimates. Latitude was standardized but not log-transformed because it includes negative values. Genus was included as a random intercept to account for potential non-independence among congeneric species. The model was fitted with 453 of the 494 valid species, due to lack of information for some species traits (e.g., body size).

**Table 1.** Biological, geographic, environmental/hydrographic, and anthropogenic-accessibility predictors used to explain variation in the year of taxonomic description of Rivulidae species.

| Variable | Unit and calculation | Justification | Source |
| --- | --- | --- | --- |
| Life-history strategy | Binary categorical: annual or non-annual | Annual Rivulidae depend on temporary habitats and can be collected only during specific periods of the hydrological cycle, whereas non-annual species can be sampled throughout the year. This difference in sampling availability may influence the timing of species discovery and description | Furness et al. (2018); Guedes et al. (2025a) |
| Range size / EOO | km<sup>2</sup>; calculated as the sum of the areas of HydroBASINS level-9 sub-basins occupied by each species | Species with broader geographic distributions are more likely to be encountered, collected, and described earlier (Moura and Jetz 2021) | HydroBASINS level 9; Lehner and Grill (2013). HydroBASINS level 9 was selected following Guedes et al. (2025a) |
| Latitude | Decimal degrees; mean latitude of the occurrence records of each species | Latitude represents the mean geographic position of each species and allows testing whether description history varied among tropical, subtropical, austral, or northern portions of the study region (Zapata and Robertson 2007) | Occurrence records compiled in this study; Guedes et al. (2025a); SALVE/ICMBio (2026a); GBIF (2026); original species descriptions |
| Elevation | Meters above sea level; mean elevation of the occurrence records of each species | Elevation may represent geographic isolation, topographic complexity, and access difficulty. Species occurring in elevated or topographically complex areas may be discovered later because these regions tend to be less accessible and have historically been less sampled (Reis et al. 2024; Guedes et al. 2025b) | SRTM/CGIAR-SRTM elevation data (Jarvis et al. 2008) |
| River order | Ordinal or numeric value representing the mean hierarchical position of species occurrence records within the drainage network | River order represents the position of each species within the hydrographic network (Lehner and Grill 2013). Species associated with headwaters, small tributaries, or lower-order reaches may be more difficult to detect than species associated with main river channels or higher-order reaches (Reis et al. 2024) | HydroBASINS level 9; Lehner and Grill (2013) |
| Population density | Inhabitants per km<sup>2</sup>; mean value across HydroBASINS level-9 sub-basins occupied by each species | Population density may act as a proxy for scientific infrastructure. More densely populated regions tend to concentrate universities, museums, biological collections, specialists, and sampling campaigns, increasing the probability of earlier discovery and description (Moura et al. 2018; Moura and Jetz 2021). | HydroBASINS level 9; Lehner and Grill (2013) |
| Road density | Meters per km<sup>2</sup>; mean value across HydroBASINS level-9 sub-basins occupied by each species | Road density is a direct proxy for accessibility. Regions with denser road networks are more easily accessed by researchers, increasing the probability of collection, deposition in biological collections, and taxonomic description (Oliveira et al. 2016; Penhacek et al. 2025) | HydroBASINS level 9; Lehner and Grill (2013) |
| Body size | Maximum total length, TL, in mm | Larger species tend to be more visible, more easily collected, and more rapidly recognized as distinct, whereas small, cryptic, or difficult-to-collect species may remain undescribed for longer periods (Moura and Jetz 2021; Reis et al. 2024) | Guedes et al. (2023a); FishBase (Froese and Pauly, 2026), via *rfishbase* (Boettiger et al. 2012); original descriptions, redescrptions, and taxonomic revisions; TL estimated from SL when necessary |

Before fitting the Bayesian model, multicollinearity among predictors was assessed using variance inflation factors (VIFs) calculated from an equivalent frequentist model, and VIF < 3 was found for all variables. Bayesian models were fitted with four Markov Chain Monte Carlo chains, each with 4,000 iterations. Posterior distributions were summarized using posterior means and 95% credible intervals. Predictors whose 95% credible intervals did not overlap zero were interpreted as having strong support for positive or negative associations with the year of species description. Model convergence was assessed using the potential scale reduction factor (R-hat), and model fit was evaluated using Bayesian R². Finally, spatial variation in the mean year of species description across the Neotropical region was mapped. Occurrence records were assigned to HydroATLAS level-9 basins, and for each basin the mean description year of all valid Rivulidae species recorded within it was calculated. A 1° grid was then overlaid across the Neotropical region, and each grid cell was assigned the mean of the basin-level description years intersecting that cell.

### 2.5. Wallacean Shortfall

To evaluate the Wallacean shortfall, spatial gaps in the geographic knowledge of Rivulidae were quantified using the cleaned occurrence dataset described above, comprising 3,419 occurrence records of 494 valid species after removing duplicated records of the same species. The Neotropical region was delimited using a bounding box, which was overlaid with a 1° grid and clipped to terrestrial areas using country polygons from the *rnaturalearth* R package (Massicotte and South 2023). Occurrence records were assigned to grid cells through spatial joins implemented in the *sf* R package (Pebesma 2018), allowing the calculation of the number of occurrence records and species recorded in each cell. To delimit the geographic occurrence envelope of Rivulidae, all occupied grid cells were merged and a concave hull was generated using the *concaveman* algorithm (Gombin et al. 2025), providing a closer approximation to the known distribution than a convex hull. Because Rivulidae are freshwater fishes frequently associated with lowland environments, high-elevation areas (> 2,000 m above sea level) were excluded from this envelope using a digital elevation model obtained with the *elevatr* package (Hollister 2025), thereby excluding regions such as the Andes from the potential geographic extent of the family (Supplementary material 1). The resulting occurrence envelope was then intersected with the terrestrial grid, and all intersecting cells were classified as either containing Rivulidae records or lacking records. The number and proportion of occupied and unoccupied cells were subsequently calculated within this occurrence envelope to quantify the proportion of unsampled areas.

To estimate sampling completeness, sample coverage for each grid cell was calculated using the *DataInfo* function from the *iNEXT* R package (Hsieh et al. 2024). For each cell, species-level record counts were treated as abundance data, and sample coverage was interpreted as the estimated proportion of the local assemblage represented by the available occurrence records. Cells without occurrence records were treated as unsampled. Following Guedes et al. (2026a), sampling coverage was used as a measure of inventory completeness, but the minimum record threshold was adapted to the biology and data availability of Rivulidae, a group characterized by small ranges and relatively few occurrence records (Guedes et al. 2025a). A grid cell was considered to be well sampled when sample coverage was at least 70% and the cell contained at least 10 occurrence records, unlike other studies that consider 50 occurrence records (Frateles et al. 2026; Guedes et al. 2026a), which is an unrealistic measure for species as cryptic and rare as Rivulidae, while 10 occurrence records demonstrates that the region was very likely revisited several times. To evaluate the sensitivity of this criterion, well-sampled cells were also mapped using alternative minimum thresholds of 5 and 20 records, while keeping the coverage threshold fixed at 70%. Finally, the geographic distance from each grid cell containing at least one Rivulidae record to the nearest well-sampled cell was calculated. Well-sampled cells were defined using the intermediate criterion of at least 70% sample coverage and at least 10 records. Distances were calculated between representative points of grid cells and expressed in kilometers. This analysis was used to identify recorded areas that remain geographically isolated from adequately sampled cells, highlighting spatial discontinuities in inventory effort across the known distribution of Rivulidae.

### 2.6. Conservation Shortfall

For all species with at least one extinction-risk assessment, the disparity between taxonomic description and conservation assessment was calculated following Tapley et al. (2018), defined as the time elapsed between the year of taxonomic description and the year of the first available assessment. Because the IUCN Red List was established in 1964, the starting year for this calculation was set to 1964 for species described before the creation of the Red List, in order to avoid artificially inflating the time without assessment. For species described after 1964, the year of taxonomic description was used as the starting point. Assessment histories were retrieved using the *rredlist* package and the *rl_species()* function (Gearty and Chamberlain 2025), which queries the IUCN Red List API for the extinction-risk assessment history associated with each species. Searches were conducted in June 2026 and included both currently valid names and all potential synonyms. This analysis was based exclusively on IUCN Red List data.

To assess the territorial protection gap, georeferenced species occurrence records were overlaid with vector layers from the World Database on Protected Areas (WDPA), obtained from Protected Planet (UNEP-WCMC and IUCN 2026). The main analysis considered only spatial records in protected areas represented by polygons, and excluded areas with “Proposed” status or with unreported designation status. Because the WDPA may contain multiple overlapping designations for the same geographic space, for example an area simultaneously recognized as a National Park, Ramsar Site, and Natural World Heritage Site, polygons were processed to avoid double counting (Deguignet et al. 2017). When polygons belonging to different protection categories overlapped the same species occurrence record, the strictest protection category was prioritized. Protected areas were grouped according to their management category, distinguishing strict protection, corresponding to IUCN categories I–IV, from non-strict or multiple-use protection, corresponding to IUCN categories V–VI (Dudley 2008; Austin et al. 2025). The overlay of occurrence data and vector layers were performed in QGIS software version 3.34 Prizren (QGIS Development Team 2026). Analyses were performed using R version 4.4.3. For code and raw data see the Data Availability.

## 3. Results

### 3.1. Linnean Shortfall

A total of 545 Rivulidae species were compiled, of which 494 were considered valid and 51 were treated as synonyms. Species description years ranged from 1821 to 2025. The annual number of valid species descriptions increased markedly after 1975, with no indication of reaching an asymptote (Fig. 2a). In contrast, the accumulation of synonyms was considerably slower and appeared to stabilize during the last decade (Fig. 2a). Consequently, the proportion of synonyms relative to valid species declined sharply around the 2000s (Fig. 2b).

**Fig. 2.**
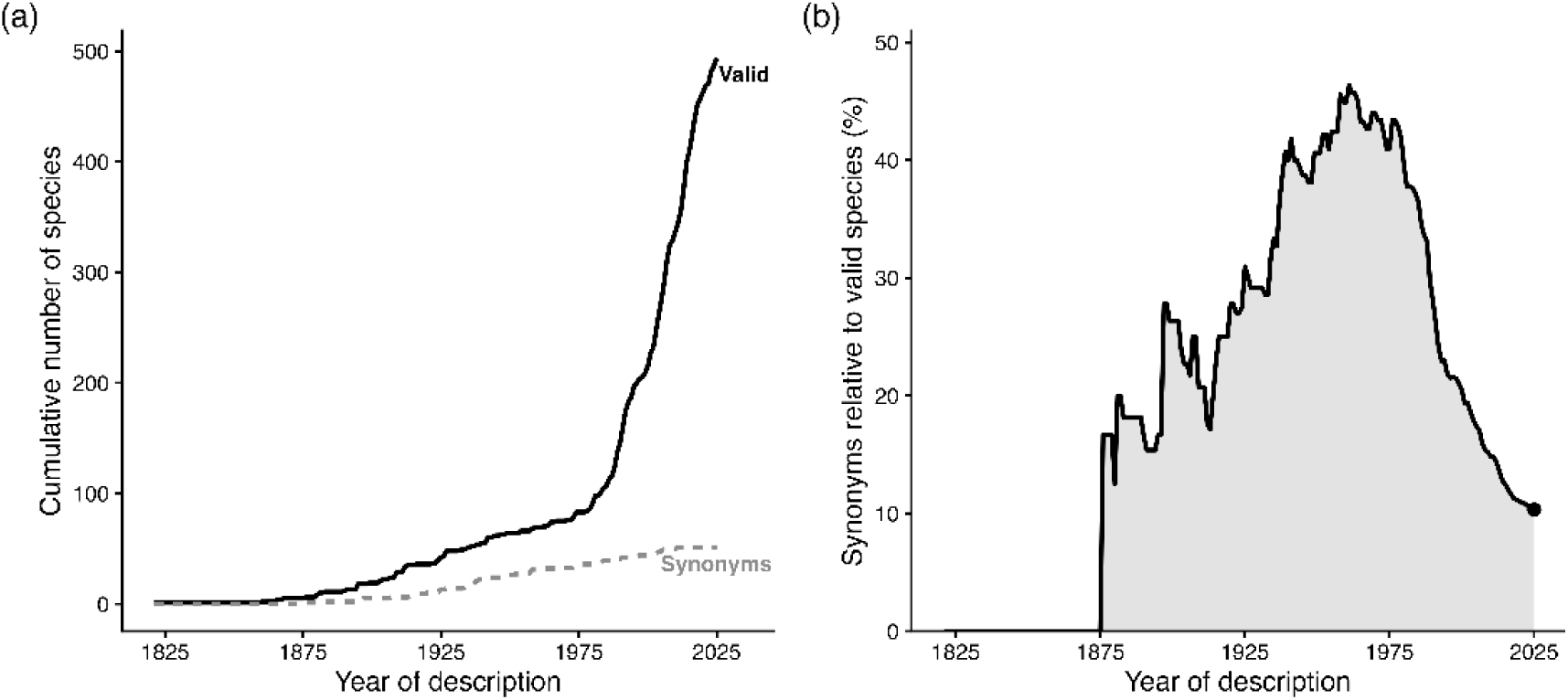
Temporal dynamics of species descriptions and synonym accumulation in Rivulidae. (a) Cumulative number of valid species (black line) and synonyms (dashed gray line) described between 1821 and 2025. (b) Temporal variation in the proportion of synonyms relative to the total number of species.

Across the entire period from 1821 to 2025, the mean description rate was 2.41 species per year for valid species. However, description rates varied substantially through time, increasing from only 0.1 species per year in the 1820s to 0.6 species per year in the 1960s, reaching 12.0 and 12.8 species per year in the 2000s and 2010s, respectively. Among the models evaluated to estimate total species richness, the Lu and He model received the strongest support (Table 2). This model predicted a total richness of 597 Rivulidae species (95% confidence interval: 547–706), suggesting that approximately 103 species remain undescribed, with estimates ranging from 53 to 212 species.

**Table 2.** Predictive species discovery models statistics for Rivulidae species described from 1821 to 2025. S_tot_ = estimated total species richness. 95% CI = 95% confidence interval. *a* and *b* = estimated model parameters. ΔAIC = Akaike Criterion Information, WAIC = AIC weights.

| Model | $S_{tot}$ | Lower<br>95% CI | Upper<br>95% CI | $a$ | $b$ | $\Delta AICc$ | $wAICc$ |
| --- | --- | --- | --- | --- | --- | --- | --- |
| Lu and He | 597 | 547 | 706 | $2.33 \times 10^{-3}$ | $7.04 \times 10^{-5}$ | 0 | 0.9978 |
| Joppa | 556 | 530 | 621 | $-3.31 \times 10^{-3}$ | $5.57 \times 10^{-5}$ | 34.5 | <0.001 |
| Logistic | 1,125 | 702 | 73,356 | $2.95 \times 10^{-4}$ | $1.49 \times 10^{-4}$ | 12.3 | 0.002 |

The Bayesian model indicated that the year of species description was associated mainly with geographic, body size, and human-associated predictors (Fig. 3). Species occurring at higher latitudes were described earlier, as indicated by the negative posterior effect of latitude (posterior mean = −0.0029, 95% credible interval (CrI): −0.0050 to −0.0007). Species with larger range sizes were also described earlier (posterior mean = −0.0036, 95% CrI: −0.0052 to −0.0019), as were larger-bodied species (posterior mean = −0.0041, 95% CrI: −0.0061 to −0.0023). Population density showed the strongest supported negative association with year of description (posterior mean = −0.0050, 95% CrI: −0.0074 to −0.0025), suggesting that species occurring in more densely populated areas tended to be described earlier. In contrast, annual life cycle, elevation, river order, and road density showed no clear association with year of description, as their 95% credible intervals overlapped zero. The model explained a moderate proportion of variation in description year (Bayesian R² = 0.29, 95% CrI: 0.22–0.36). Variation among genera was negligible, with the random intercept for genus showing an estimated standard deviation close to zero (SD = 0.00, 95% CrI: 0.00–0.01), indicating that description patterns were largely independent of genus-level taxonomic structure. Convergence diagnostics indicated reliable sampling, with a maximum R-hat of 1.002.

**Fig. 3.**
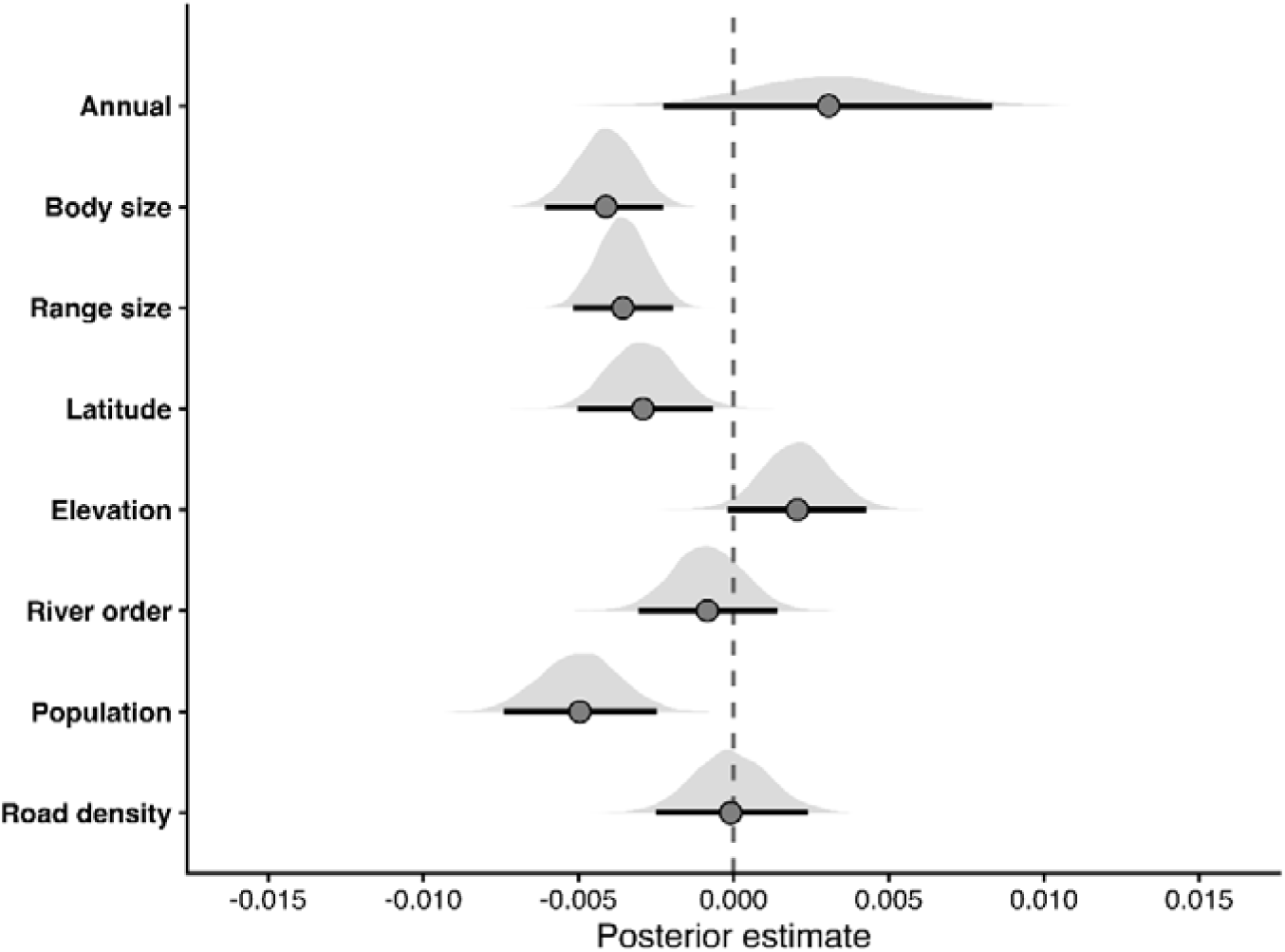
Predictors of species description year in Rivulidae. Posterior distributions of fixed effects from a Bayesian Gamma model evaluating predictors of the year of species description. Points indicate posterior means and horizontal bars represent 95% credible intervals. The dashed vertical line indicates zero effect. Negative estimates indicate predictors associated with earlier species descriptions, whereas positive estimates indicate predictors associated with more recent descriptions. Effects whose 95% credible intervals do not overlap zero are interpreted as supported predictors.

Spatial patterns in the mean year of species description revealed marked geographic variation across the Neotropical region (Fig. 4a). Cells located in Central America, the Caribbean, and northern portions of South America generally contained species described earlier, whereas many other areas in South America were characterized by more recent average description years. Particularly high mean description years were concentrated in Bolivia, and in eastern and central Brazil, as well as in several regions of the Amazon Basin, indicating that these areas harbor a greater proportion of recently described species (Fig. 4a).

**Fig. 4.**
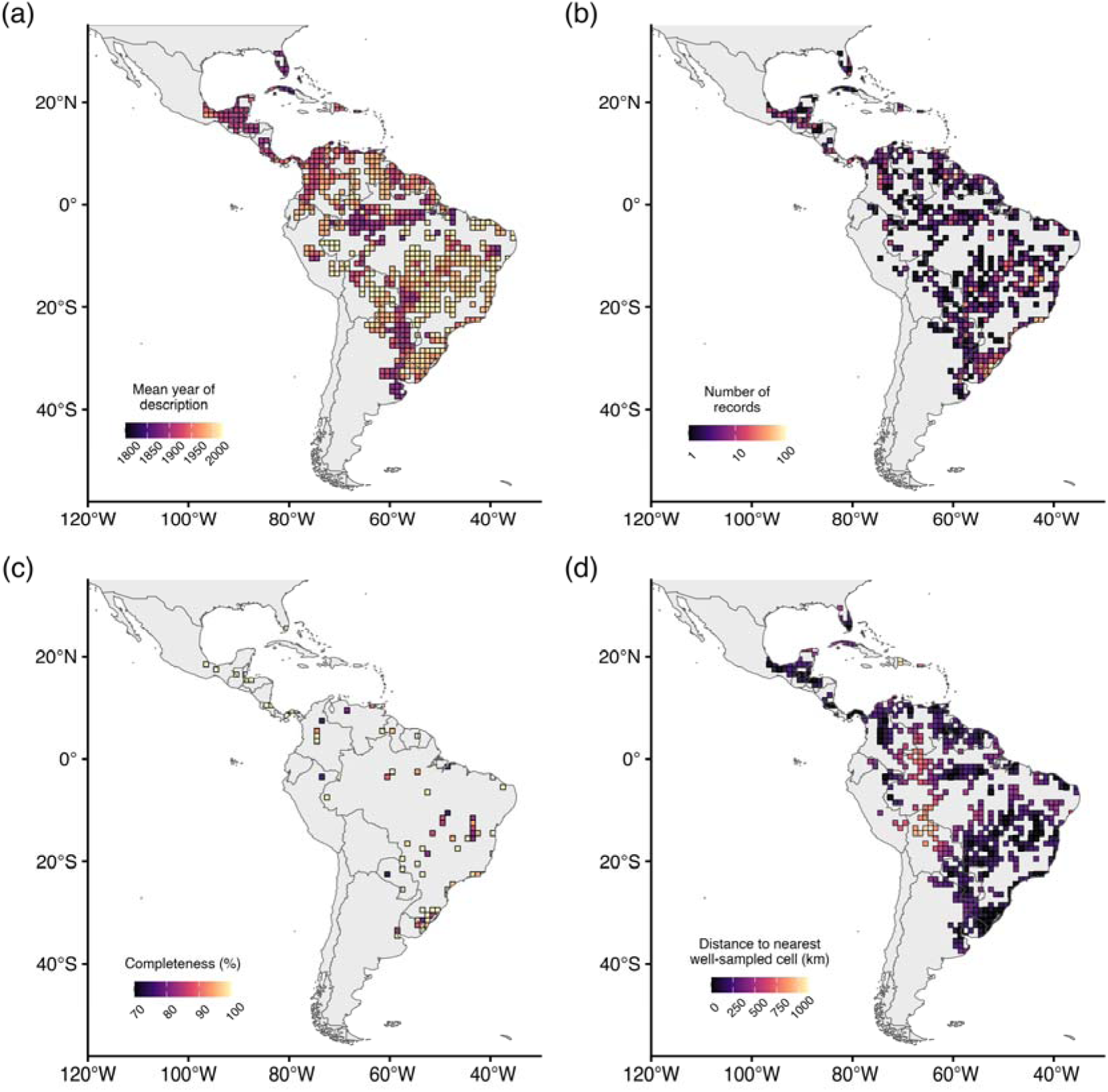
(a) Spatial variation in the mean year of species description across the Neotropical region. Values represent the mean description year of Rivulidae species within each 1° grid cell. Darker colors indicate cells dominated by species described earlier, whereas lighter colors indicate cells containing a greater proportion of recently described species. (b) Spatial distribution of Rivulidae occurrence records per 1° grid cell. (c) Sampling completeness estimated using sample coverage, considering cells with at least 70% coverage and minimum record threshold of 10 occurrence records. Cells not meeting these criteria are not shown. (d) Geographic distance from each Neotropical grid cell containing Rivulidae records to the nearest well-sampled cell. Well-sampled cells were defined as those with at least 70% sampling coverage and a minimum of 10 occurrence records. Warmer colors indicate greater distances to the nearest adequately sampled area, highlighting regions where occurrence data remain geographically isolated from well-inventoried cells.

### 3.2. Wallacean Shortfall

Based on 3,419 unique Rivulidae occurrence records, the occurrence envelope comprised 1,536 terrestrial 1° grid cells, of which 950 (61.85%) lacked any occurrence records, indicating that most of the potential geographic extent of the family remains unsampled. Among the 586 cells containing records, sampling effort was highly uneven, ranging from 1 to 120 records per cell (mean = 5.6; median = 3), with cells in Uruguay and southern Brazil presenting the highest numbers of records (Fig. 4b). Species richness per occupied grid cell varied from 1 to 15 species, with a mean of 2.04 and a median of one species per cell. When applying a minimum sampling coverage threshold of 70%, combined with minimum record requirements of 5, 10, or 20 records per cell, only 161, 82, and 33 cells, respectively, were classified as well-sampled (Fig. 4c; Supplementary material 2). Using the intermediate criterion of at least 10 records and 70% sampling coverage (Fig. 4c), only 5.34% of the grid cells within the Rivulidae occurrence envelope were considered adequately sampled, being concentrated mainly in southern and southeastern Brazil, with additional well-sampled cells in central Brazil, Colombia, and Central America. The spatial distribution of distances to the nearest well-sampled cell revealed pronounced geographic disparities in sampling coverage across the occurrence envelope (Fig. 4d). Cells classified as well-sampled exhibited distances of zero by definition, whereas many cells containing Rivulidae records were located hundreds of kilometers from the nearest adequately sampled area. The largest distances were concentrated in central South America, particularly in portions of Bolivia and the Brazilian Amazon (Fig. 4d).

### 3.3. Conservation Shortfall

Among the 494 species analyzed, 425 (86%) had undergone some assessment of extinction risk (Fig. 5), whereas 69 (14%) had not yet been evaluated (NE). Within the subset of evaluated species, 179 (42.2%) were classified as threatened with extinction (CR, EN, or VU), 184 (43.2%) as non-threatened (NT or LC), and 62 (14.5%) as Data Deficient (DD). Species with an annual life cycle disproportionately dominated the threatened fraction of Rivulidae, comprising 80.4% (144 of 179) of all species classified as threatened. Ten (10) species were identified as Possibly Extinct (PE), representing a subset of CR species for which the available evidence suggests probable extinction, although confirmation remains insufficient for formal classification as Extinct (EX). These included *Cynolebias elegans* Costa, 2017, *Hypsolebias splendissimus* Costa, 2018, *Matilebias toba* (Calviño, 2006), *Moema claudiae* (Costa, 2003), and *Ophthalmolebias perpendicularis* (Costa, Nielsen & de Luca, 2001), identified as PE in the IUCN Red List; and *Atlantirivulus nudiventris* (Costa & Brasil, 1991), *Hypsolebias marginatus* (Costa & Brasil, 1996), *Leptopanchax sanguineus* Costa, 2019, *Moema piriana* Costa, 1989, and *Simpsonichthys zonatus* (Costa & Brasil, 1990), identified as PE by SALVE/ICMBio.

**Fig. 5.**
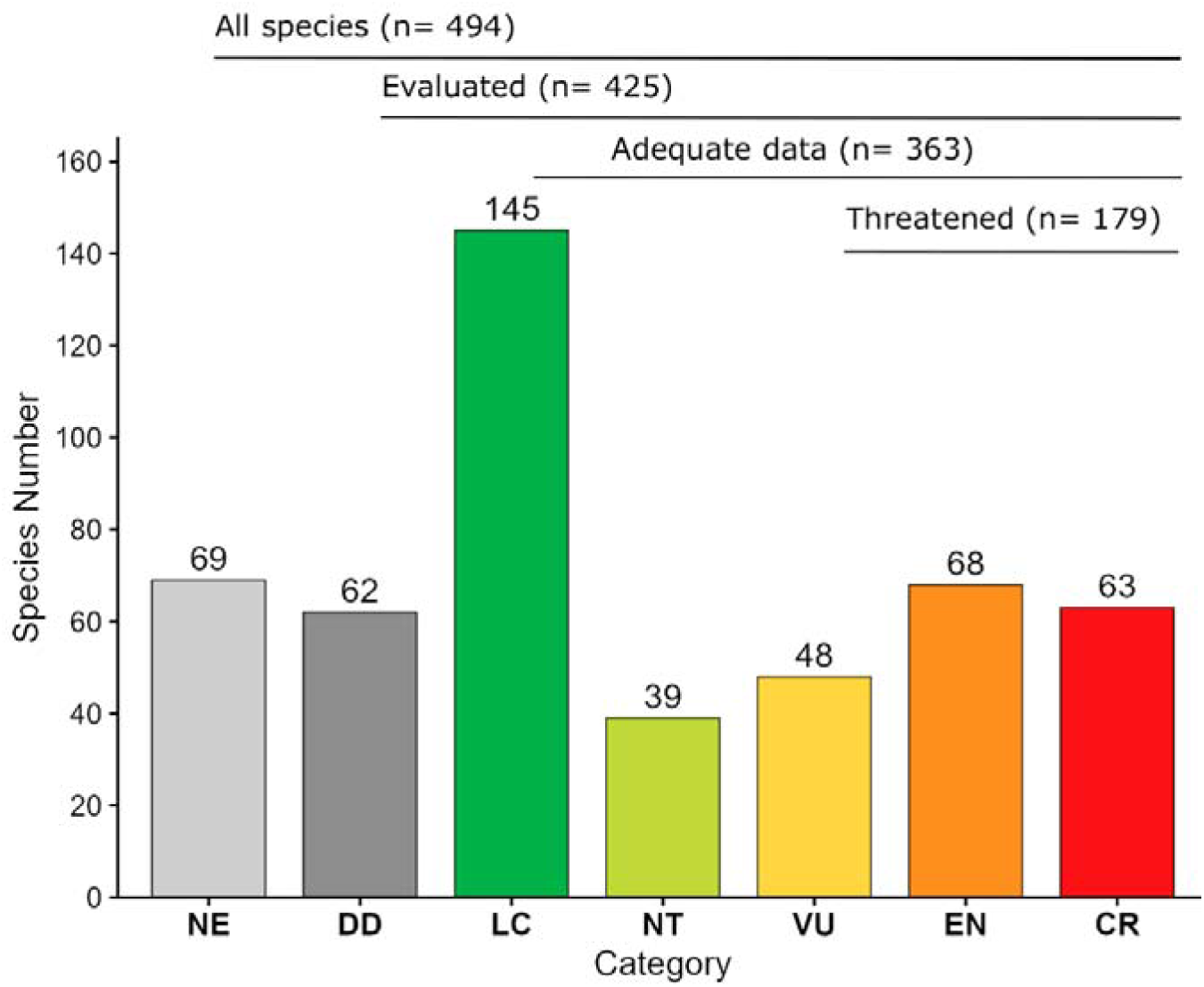
Number of Rivulidae species in each IUCN conservation category, combining data from the IUCN Red List and SALVE/ICMBio.

The temporal gap between taxonomic description and formal conservation assessment was substantial in Rivulidae. After accounting for the establishment of the IUCN Red List in 1964, the mean time to first assessment was 25.7 years (median = 21 years), ranging from 2 to 60 years. The first Rivulidae assessments were not conducted until 1986, when eight (8) species were incorporated into the Red List (e.g., *Leptolebias marmoratus* (Ladiges, 1934), *Millerichthys robustus* (Miller & Hubbs, 1974), and *Ophthalmolebias constanciae* (Myers, 1942)), despite 113 species already being taxonomically described and available for assessment at that time (Figure 6). Annual species accounted for the majority of assessed taxa (231 species; 58.8%) and exhibited a slightly shorter mean time to first assessment (24.8 years; median = 21 years) than non-annual species (162 species; 41.2%; mean = 26.9 years; median = 23.5 years). Assessment delays were spatially heterogeneous: species occurring in Brazil showed a shorter mean time to first assessment (23.1 years; median = 19 years) than species not occurring in Brazil (32.1 years; median = 29 years). Among countries with the largest numbers of assessed species, the longest mean delays were observed for species occurring in Colombia (42.8 years), Argentina (42.1 years), Uruguay (37.5 years), Panama (36.4 years), Venezuela (34.1 years), Bolivia (33.3 years), and Peru (32.5 years). The incorporation of species into the IUCN Red List was strongly concentrated in recent years: 315 species, representing 80.2% of taxa with available assessment data, received their first assessment between 2021 and 2024, with a pronounced peak in 2022, when 170 species were assessed for the first time (Fig. 6).

**Fig. 6.**
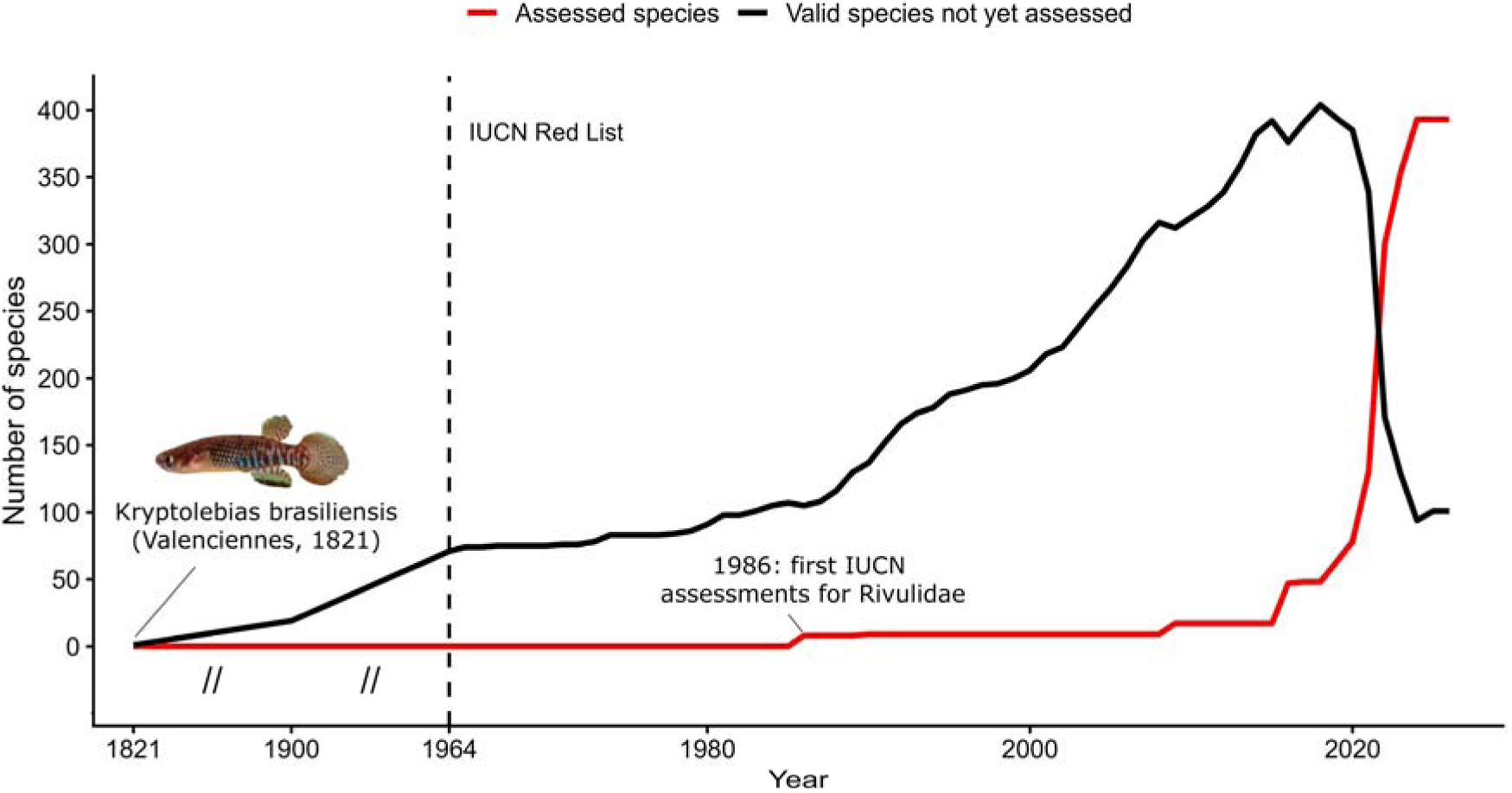
Temporal dynamics of the conservation assessment shortfall in Rivulidae. The black line represents the number of valid species that were taxonomically available but had not yet been assessed by the IUCN Red List in each year. The red line represents the cumulative number of species formally assessed by the IUCN. The dashed vertical line marks the establishment of the IUCN Red List in 1964. The broken x-axis highlights the early taxonomic history of Rivulidae, marking 1821 as the year in which the first species of the family was described, *Kryptolebias brasiliensis* (Valenciennes, 1821), while preserving the temporal dynamics following the establishment of the IUCN Red List.

Regarding territorial protection gaps, 296 species (59.9%) had no records within WDPA protected areas, whereas 198 species (40.1%) had at least one record overlapping protected areas. Considering mutually exclusive groups, 39 species (7.9%) occurred in strict protection areas, 107 species (21.7%) occurred in sustainable-use protected areas, and 52 species (10.5%) occurred simultaneously in both strict and sustainable-use areas. Regarding occurrence records, 755 of the 3,419 records analyzed (22.1%) overlapped protected areas. Of these, 251 records (7.3% of the total) occurred in strict protection areas and 504 records (14.7%) in non-strict or sustainable-use protected areas. Most occurrence records of rivulid fishes (78.2%; 2,674 records) were located in unprotected areas. Among the 179 threatened species (VU, EN, CR), only 55 (30.7%) had at least one distribution record overlapping protected areas.

## 4. Discussion

### 4.1 Linnean Shortfall

Overall, we found a substantial Linnean shortfall for Rivulidae fishes. The cumulative curve of valid species descriptions has not yet reached an asymptote, indicating that a considerable fraction of the family’s diversity likely remains undescribed. Similar patterns have been reported for other vertebrates (Frateles et al. 2024), including fishes (Freitas et al. 2021). However, the absence of an asymptote may also reflect the high rate of species descriptions during the last two decades, which could be associated with increased research capacity and scientific interest in the group. In this context, although the number of practicing taxonomists has declined globally (Pearson et al. 2011), taxonomic effort still appears to be a major driver of research focus in Rivulidae and in the freshwater environments they typically inhabit (Costa et al. 2025). At the same time, the number of synonyms recorded for the family was relatively low (51 species), and the proportion of synonyms relative to valid species declined markedly after 2000. This pattern is likely explained by two complementary processes. First, the accelerated description of valid species in recent decades dilutes the relative contribution of synonyms to the total species names. Second, species descriptions have become increasingly robust, following a trend observed in other vertebrate groups (Guedes et al. 2024), thereby reducing the likelihood that newly described taxa will later be synonymized. Indeed, many recent Rivulidae descriptions integrate molecular evidence with traditional morphological diagnoses (Abrantes et al. 2023; Volcan et al. 2025), and often include complementary information on ecology, distribution, and environmental threats (Alonso et al. 2024, 2025; Abrantes et al. 2026). This pattern is particularly relevant because taxonomic uncertainty can directly influence estimates of biodiversity patterns and conservation planning (Meyer et al. 2026). Nevertheless, the determinants of taxonomic uncertainty and the factors affecting the probability of synonymization remain poorly understood in Rivulidae and represent promising avenues for future research.

Regarding species discovery models, the Lu and He model estimated that approximately 103 Rivulidae species remain undescribed, although uncertainty was substantial, with estimates ranging from 23 to 212 species. This number may appear modest when compared with hyperdiverse groups such as Brazilian butterflies, for which more than 600 species are still expected to be described (Guedes et al. 2026a), yet it is considerably higher than estimates for New World coral snakes, where approximately 40 species remain undiscovered (Frateles et al. 2024). Comparisons with other fish groups provide more useful context; for example, approximately 45% of the expected diversity of Auchenipteridae catfishes remains undescribed (Freitas et al. 2021), whereas our estimates indicate that, on average, approximately 17.25% of Rivulidae diversity is still awaiting formal description. Although proportionally lower, this value should be interpreted in light of the biological and taxonomic context of the family. Rivulidae has experienced an exceptional taxonomic effort during recent decades (Costa et al. 2025), resulting in a remarkable increase in the rate of species descriptions. Consequently, an estimate of more than one hundred undescribed species represents a substantial challenge despite this sustained taxonomic activity. Moreover, the characteristics of recently described species suggest that most of the remaining undescribed diversity is increasingly concentrated in species that are small-bodied, have restricted geographic ranges, and occur in remote areas with low human population density, making them inherently more difficult to detect and describe. As with any species discovery model, however, these estimates remain associated with considerable uncertainty (Frateles et al. 2024; Faurby and Farooq 2026). Indeed, the logistic model produced biologically unrealistic estimates in our analyses. Nevertheless, modern discovery models that explicitly account for temporal variation in taxonomic effort by incorporating the number of active taxonomists through time (Joppa et al. 2011; Lu and He 2017; Frateles et al. 2024), as was the case for our best-performing model, provide a more realistic framework for estimating undocumented diversity. Despite their limitations, these estimates remain essential for guiding research priorities, directing funding efforts, and increasing societal awareness of the urgent need to document biodiversity before additional species are lost.

The association between more recent description years and both smaller range sizes and smaller body sizes is also consistent with patterns reported for other taxonomic groups (Frateles et al. 2024). What is particularly noteworthy is that these relationships remain evident even within Rivulidae, a family already characterized by generally small body sizes and restricted geographic distributions (Guedes et al. 2026b). Despite the relatively limited variation in these traits, with only a few species representing extreme values (e.g., Alonso et al. 2025), both body size and range size emerged as important predictors of species description timing. This pattern is likely explained by the fact that larger-bodied species with broader distributions are generally easier to detect, collect, and distinguish from related taxa (Guerra et al. 2020). Conversely, species that remain undescribed are expected to be smaller, geographically restricted, and more difficult to encounter in the field.

We also found that species occurring at higher latitudes and in more densely populated regions tended to have been described earlier. This pattern likely reflects historical and contemporary biases in sampling effort, accessibility, proximity to research institutions, and availability of funding, all of which are generally greater in more populated regions and at higher latitudes within the Neotropics (Moura et al. 2018; Moura and Jetz 2021). Together, these results suggest that future discoveries are most likely to involve species with small body sizes, restricted distributions, and occurrences concentrated in low-latitude and sparsely populated regions. The spatial distribution of mean description years strongly supports these predictions. Areas characterized by more recent average description dates were concentrated in northeastern Brazil, where several species have been described in recent years (e.g., Abrantes et al. 2026), as well as in central, southeastern, and southern Brazil (e.g., Volcan et al. 2018, 2025), the Brazilian Amazon (e.g., Nielsen et al. 2025), and portions of Bolivia (e.g., Drawert 2022). These regions appear to represent contemporary hotspots of taxonomic discovery and therefore constitute promising targets for future field surveys and taxonomic research aimed at reducing the Linnean shortfall in Rivulidae.

### 4.2 Wallacean Shortfall

When assessing the Wallacean shortfall in Rivulidae, we found extensive gaps in geographic knowledge across the family’s potential distribution, indicating that the distribution of many species remains poorly documented. Using a concave occurrence envelope encompassing all occupied grid cells while excluding high-elevation areas (>2,000 m above sea level), 950 of the 1,536 grid cells (61.85%) within the potential distribution lacked any Rivulidae record. Among the 586 occupied cells, sampling effort was highly heterogeneous, ranging from 1 to 120 records per cell, with the highest concentrations occurring in southern Brazil and Uruguay. Similar spatial biases have been reported for other taxa and are frequently associated with accessibility, proximity to research institutions, biological collections, and long-term scientific investment (Guedes et al. 2026a). Furthermore, only 82 grid cells met our criteria for being considered well-sampled, representing just 5.34% of the potential occurrence envelope. This pattern likely reflects the strong historical emphasis of Rivulidae research on taxonomy, which remains essential given the urgent need to document species before ongoing environmental changes drive further biodiversity loss. As a consequence, comparatively fewer studies have focused on assemblage-level ecology or long-term monitoring (Costa et al. 2025), research approaches that typically provide broader spatial coverage and repeated sampling of the same localities.

The spatial distribution of well-sampled cells further highlights important geographic disparities in sampling effort. Similar to patterns reported for other vertebrate groups (Frateles et al. 2026), well-sampled cells were concentrated near regions with strong scientific infrastructure. In the case of Rivulidae, these cells were overwhelmingly clustered in southern Brazil and Uruguay, consistent with the high scientific productivity of some Rivulidae-focused research groups in these regions (Costa et al. 2025). In contrast, cells located farthest from any well-sampled area were concentrated primarily in the Brazilian Amazon and parts of Bolivia. In Bolivia, this pattern likely reflects the relatively recent but growing efforts of local specialists to document Rivulidae diversity (Drawert 2022, 2023; Drawert and Ergueta 2024). In the Brazilian Amazon, however, it may indicate that, despite ongoing species discoveries and taxonomic advances (Nielsen et al. 2025), occurrence data remain insufficient to achieve adequate sampling coverage. This is particularly expected given that research in Amazonian wetlands is strongly constrained by accessibility and distance from biodiversity institutions (Carvalho et al. 2023), factors that continue to limit progress in reducing the Wallacean shortfall across remote regions.

Even so, by identifying the major geographic gaps in current knowledge, our results provide an initial step for prioritizing future surveys in regions with low record density and poor sampling completeness. Although important compilations of occurrence records have previously been conducted for individual basins, regions, and biomes (Abrantes et al. 2020; Sarmento-Soares et al. 2026), this is the first study to assemble a comprehensive continental-scale dataset to investigate large-scale patterns of biodiversity knowledge gaps across the family. Such prioritization is particularly important given funding constraints and may represent an important first step toward reducing the Wallacean shortfall in one of the most threatened groups of Neotropical freshwater fishes. Ultimately, species cannot be effectively conserved where they remain undocumented, making the reduction of the Wallacean shortfall a fundamental prerequisite for the long-term conservation of Rivulidae diversity.

### 4.3 Conservation Shortfall

With 179 threatened species, Rivulidae ranks fourth among fish families in absolute number of threatened species, after Cyprinidae (370 spp.), Cichlidae (314 spp.), and Leuciscidae (195 spp.) (IUCN 2026). However, relative to its diversity, Rivulidae has the highest proportion of threatened species among these highly diverse families: 36.2% of its 494 valid species are threatened, exceeding Leuciscidae (27.3% of 713 spp.), Cyprinidae (20.4% of 1,816 spp.), and Cichlidae (17.8% of 1,768 spp.) (Fricke et al. 2026b; IUCN 2026). This exceptional level of threat may still be underestimated, as 131 Rivulidae species remain either Not Evaluated (NE) or Data Deficient (DD), categories that may conceal high extinction risk (Borgelt et al. 2022). Annual species accounted for 80.4% of all threatened Rivulidae, highlighting the disproportionate vulnerability of taxa dependent on temporary wetlands. The main threat to annual species is land-use and land-cover change in temporary wetlands, leading to habitat loss (Costa 2012; Volcan and Lanés 2018; Guedes et al. 2023b). These habitats are being systematically converted into croplands, drained and filled for pastures, urban expansion, and industrial development (Calhoun et al. 2017; Castro and Polaz 2020). Additional impacts, including fire during the dry phase of temporary wetlands (Guedes and Araújo 2026), plastic pollution (Guedes et al. 2026b), and chemical pollution associated with the widespread use of agrochemicals (Godoy et al. 2025a), further compound the pressure on an already severely threatened fauna.

The mean delay of 25.7 years between taxonomic description and the first IUCN extinction-risk assessment for Rivulidae, reveals an important temporal disconnect between the scientific recognition of species and their incorporation into formal conservation instruments. The IUCN Red List is central to translating biodiversity knowledge into conservation priorities and policies (Betts et al. 2020), but its effectiveness depends on regular reassessments and on the speed with which described species are evaluated (Tapley et al. 2018). When this process takes decades, extinction-risk assessment may represent a delayed diagnosis of populations and habitats that have already been severely reduced. Historically, the IUCN Red List has been affected by strong taxonomic and regional disparities, and freshwater fishes, especially in megadiverse regions such as South America, have remained comparatively underassessed (Miqueleiz et al. 2020). This scenario has only recently begun to change. The peak in Rivulidae assessments between 2021 and 2024 does not indicate the recent emergence of conservation concern for the group, but rather a delayed correction of this historical gap, driven by the integration of regional assessment data, particularly from Brazil (Castro and Polaz 2020; ICMBio 2026a), with broader global efforts to complete IUCN Red List assessments for freshwater fauna (Sayer et al. 2025).

The conservation shortfall in Rivulidae also extends to effective territorial protection. Although 40.1% of species had at least one record overlapping protected areas, most species (59.9%) and occurrence records (78.2%) remain outside the WDPA protected-area network. This pattern is consistent with global evidence showing that protected-area networks still have major representativeness gaps and are often established in areas less exposed to land conversion, such as remote, steep, or low-agricultural-suitability regions (Rodrigues et al. 2004; Joppa and Pfaff 2009). For Rivulidae, this limitation is particularly critical among threatened species, of which only 30.7% had any record within protected areas, indicating that formal recognition of extinction risk has not been sufficiently translated into territorial protection. Moreover, the legal designation of a protected area alone does not guarantee the containment of all anthropogenic pressures (Geldmann et al. 2019; Li et al. 2024). For example, the establishment of protected areas was effective in slowing land-use and land-cover conversion and promoting the recovery of natural matrices in habitats of the annual fish *Notholebias minimus* (Myers, 1942) (Guedes et al. 2023b). In contrast, protected areas did not consistently reduce fire frequency in temporary wetlands occupied by Brazilian annual species, probably because fires often originate outside reserve boundaries and spread into these areas without effective landscape-scale control (Guedes and Araújo 2026).

#### 4.3.1 Conservation initiatives

Despite this severe scenario of extinction risk, conservation initiatives for Rivulidae should be recognized as efforts that shift the focus from diagnosis to action. Brazil stands out for having a public instrument specifically directed at Rivulidae: The National Action Plan for the Conservation of Threatened Rivulid Fishes, coordinated since 2013 by the National Center for Research and Conservation of Continental Aquatic Biodiversity (CEPTA/ICMBio). This action plan includes 159 target species, of which 130 are nationally threatened with extinction, and brings together actions related to research, conservation, monitoring, and institutional coordination (ICMBio 2025a). One of its main practical outcomes was the development of protocols to guide environmental licensing in areas where threatened rivulids occur, strengthening the incorporation of the group into public conservation policies (ICMBio 2025b). In the context of scientific outreach and environmental education on social media, initiatives such as @peixesdacaatinga and @killifish_foundation stand out as platforms dedicated to publicizing species, habitats, and conservation actions. A visible example of this type of outreach was the description of *Hypsolebias lulai* Ramos, Nielsen, Abrantes, Lira & Lustosa-Costa, 2023, a species whose discovery was triggered by citizen science and by expeditions funded through crowdfunding (Ramos et al. 2023). *Ex situ* management initiatives and protocols have also begun to be developed, including integrated conservation approaches (ICMBio 2026b) and topsoil translocation strategies involving diapausing egg banks (Godoy et al. 2025b).

Although there is relative consensus on the importance of formal instruments for conserving Rivulidae, the role of aquarists remains marked by strong divergence. In Brazil, for example, the collection, maintenance, commercialization, and export of threatened species for ornamental purposes are strongly restricted or prohibited (SAP/MAPA Ordinance No. 17/2021). Nevertheless, threatened Brazilian rivulids are currently maintained in aquariums in different regions of the world. From a legalistic perspective, without traceable information on the origin, legal status, and continuity of aquarium-maintained populations, it is difficult to determine whether *ex situ* maintenance reduces pressure on wild populations or depends on the recurrent removal of individuals from nature, which may increase extinction risk (Borges et al. 2021, 2022). On the other hand, it is impossible to understand the taxonomic and conservation history of Rivulidae without recognizing the role of aquarists, naturalists, and non-academic collaborators. Many species known today were initially located, maintained, photographed, or publicized by these informal networks, which often reached populations and habitats before institutional science (Costa 2009). Initiatives such as the International Rivulid Conservation Program, established in 2021, seek to organize this experience into a coordinated network of *ex situ* breeding and, in 2025, included 286 Rivulidae populations maintained in aquariums worldwide (rivulid-conservation.org). This network is particularly relevant because it includes species of exceptional conservation concern, such as *Simpsonichthys zonatus* (Costa & Brasil, 1990), which is maintained in aquaria and has been considered possibly extinct (Pavanelli et al. 2024b). Beyond legal controversies, this initiative shows that aquarists have consolidated an international network of knowledge and *ex situ* maintenance at a time when formal conservation structures were still advancing slowly. Therefore, if the ultimate goal is conservation, the debate cannot be reduced to a simple opposition between legality and hobby. Different perspectives need to sit at the same table to share the responsibility of building transparent, legally secure, and biologically responsible protocols.

### 4.4 Limitations, implications and future directions

Although this study provides valuable insights, some inherent limitations should be acknowledged. First, estimates of valid species, synonyms, threat status, and geographic distribution are inherently dynamic. Therefore, our study should be interpreted as a temporal snapshot of the current state of knowledge, although it represents the most comprehensive synthesis yet produced for Rivulidae knowledge gaps. Second, the estimated number of undescribed species should be interpreted with caution because species discovery models rely on historical description rates and are sensitive to assumptions about taxonomic effort, description year, methodological change, and future taxonomic revisions (Joppa et al. 2011; Frateles et al. 2024; Faurby and Farooq 2026). Third, species description patterns in Rivulidae are strongly influenced by the historical concentration of taxonomic effort. A single taxonomist, Wilson J. E. M. Costa, is responsible for the description of 233 species (47%) of the currently valid species of Rivulidae, and without his work, the knowledge gap addressed in this study would be substantially larger. However, this individual effect is difficult to incorporate quantitatively into the model, as it reflects individual expertise, access to collections, field experience, and long-term engagement with the group. Consequently, some associations between biological or geographic predictors and year of description may partly capture the spatial and temporal footprint of specialist activity, rather than intrinsic differences in species detectability alone. Overall, the Linnean shortfall shows that conservation strategies operate on underestimated biodiversity, in which species that remain undescribed may already be threatened or, as in *Leptopanchax sanguineus* Costa, 2019, may be formally recognized only when possibly extinct (Costa 2019).

The Wallacean shortfall indicates that spatial conservation decisions are still based on an incomplete geography. Despite our compilation of distributional data, many countries within the natural distribution of Rivulidae were formerly colonies of Global North nations, and numerous specimens collected in the Global South remain deposited in foreign institutions, where historical collections have not yet been fully digitized (Nakamura et al. 2025). In addition, challenges associated with species identification, data storage, and biodiversity data mobilization in countries with relatively few specialists continue to constrain the availability of occurrence records across much of the Global South (Frateles et al. 2026), despite important initiatives such as the Brazilian SALVE system (ICMBio 2026a). In a complementary way, the conservation shortfall exposes that describing a species does not guarantee its short-term assessment, assessing it as threatened does not guarantee territorial protection, and protecting it solely through the legal designation of protected areas may not be sufficient to ensure species persistence.

Future research should prioritize inventories in undersampled regions, especially Bolivia, the Brazilian Amazon, and central portions of South America, where records remain distant from the best-sampled areas. Taxonomic revisions integrating morphology, genetics, distribution, and natural history are also needed to resolve species complexes with restricted distributions and reduce taxonomic uncertainties. In parallel, Data Deficient and recently described species should be urgently assessed or reassessed, particularly those associated with temporary wetlands undergoing rapid conversion. In applied conservation, it is necessary to expand the spatial coverage of protected areas, while also developing complementary *in situ* and *ex situ* conservation strategies outside these areas, including private properties, environmental education, and licensing instruments. Ultimately, despite considerable knowledge shortfalls, the conservation of Rivulidae cannot wait for complete knowledge.

## Acknowledgments

We acknowledge all researchers, institutions, and professionals working in favor of rivulid conservation, with special recognition to the Centro Nacional de Pesquisa e Conservação da Biodiversidade Aquática Continental — CEPTA/ICMBio, for its efforts in the assessment and conservation of Brazilian continental aquatic biodiversity.

## Data availability

The step-by-step analysis is available in R code, along with the raw data, in a GitHub repository (https://github.com/JH-All/Rivulidae_shortfalls).

## Contributions

Conceptualization: J.H.A.C., G.H.S.G.; Investigation: J.H.A.C., G.H.S.G.; Methodology: J.H.A.C., G.H.S.G.; Formal analysis: J.H.A.C., G.H.S.G.; Data curation: J.H.A.C., G.H.S.G.; Funding acquisition: J.H.A.C., G.H.S.G.; Resources: J.H.A.C., G.H.S.G.; Visualization: J.H.A.C., G.H.S.G.; Validation: J.H.A.C., G.H.S.G.; Writing – original draft: J.H.A.C., G.H.S.G.; Writing – review & editing: J.H.A.C., G.H.S.G.

## SUPPLEMENTARY MATERIAL

**Figure S1.**
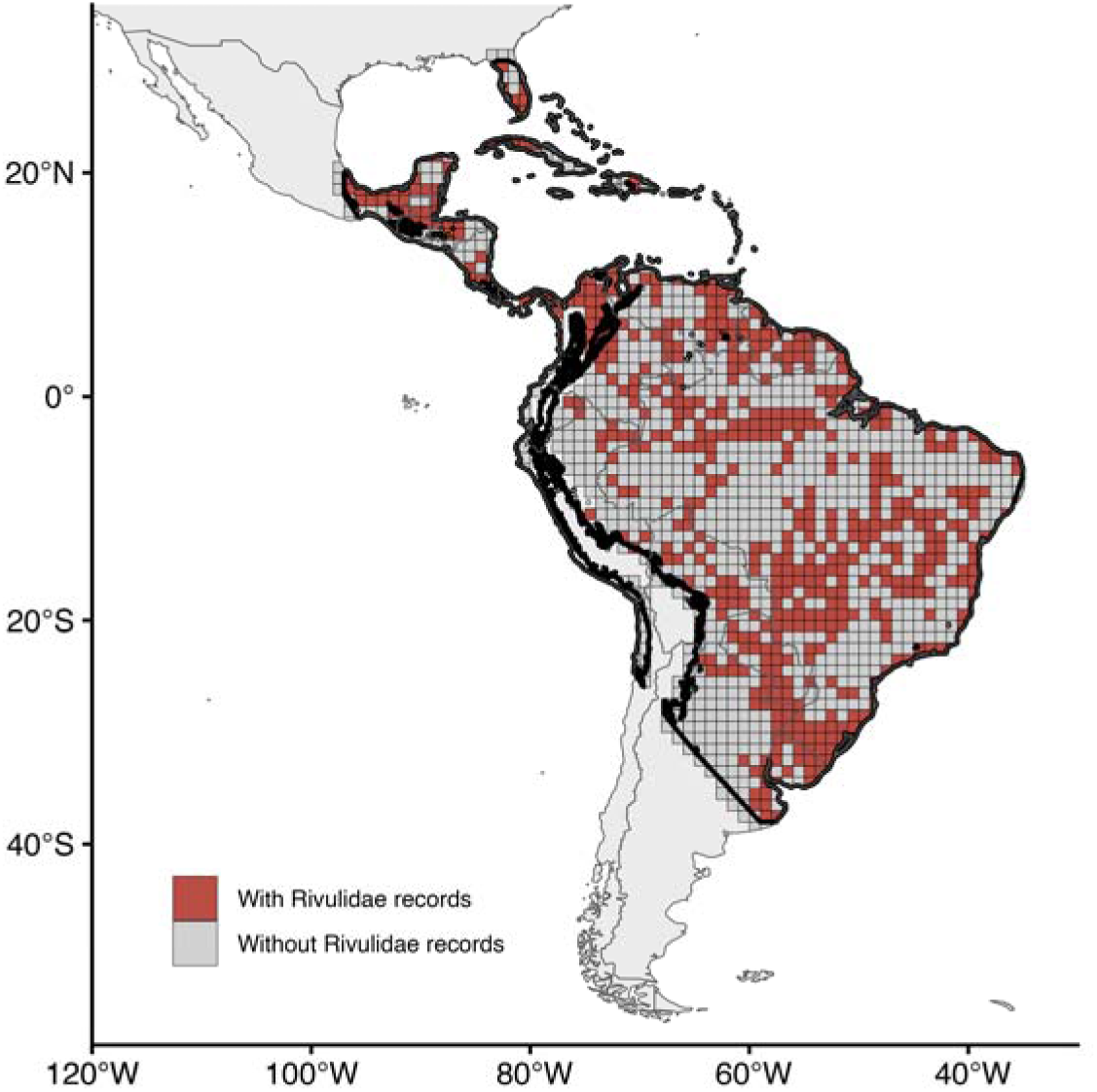
Delimitation of the Rivulidae occurrence envelope used to quantify the Wallacean shortfall. The envelope was generated by applying a concave hull to all occupied 1° grid cells and subsequently excluding high-elevation areas (> 2,000 m above sea level) to avoid including regions with low occurrence potential, such as the Andes. Grid cells within the resulting occurrence envelope are classified as containing Rivulidae records (red) or lacking records (gray).

**Figure S2.**
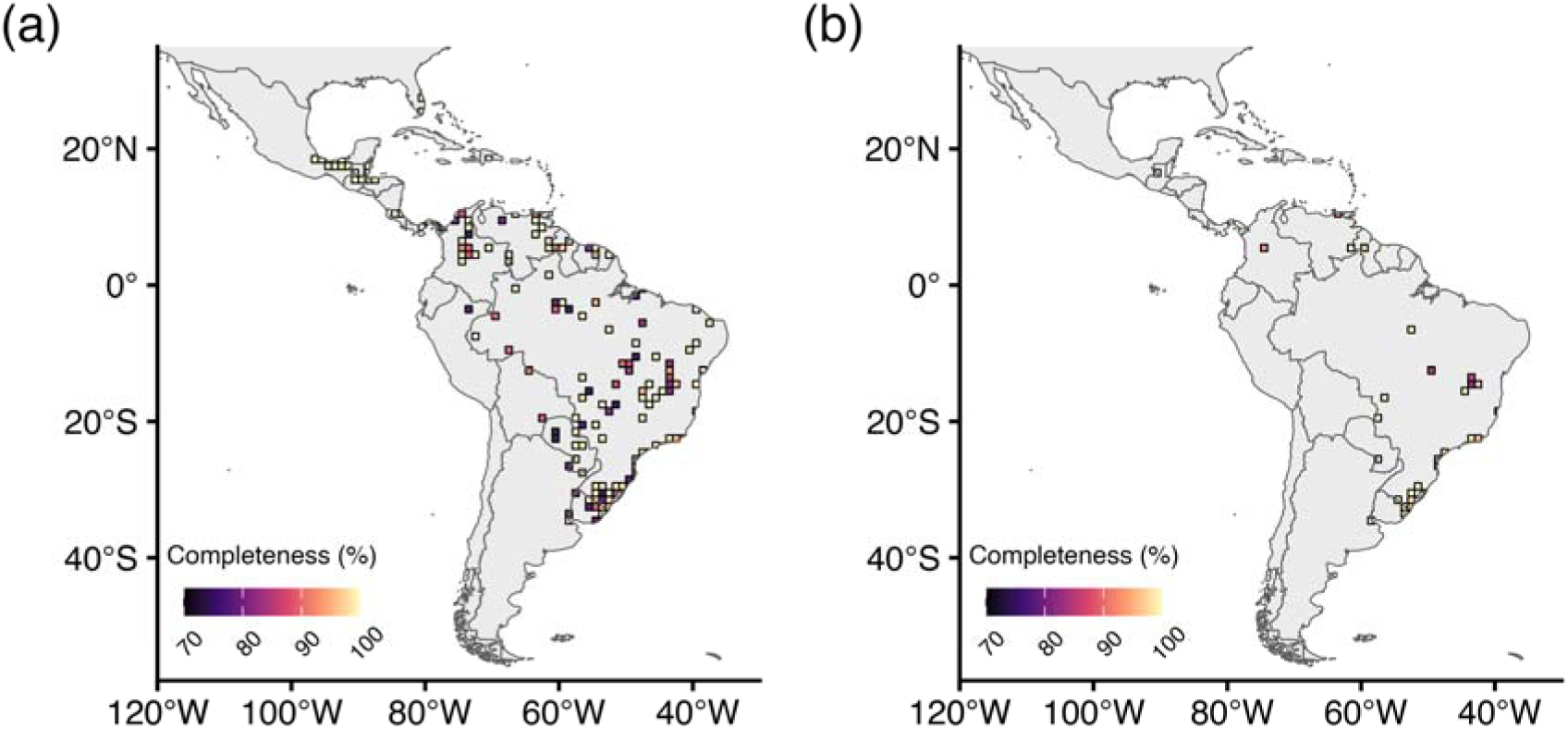
Sampling completeness for Rivulidae estimated using sample coverage, considering cells with at least 70% coverage and minimum record thresholds of 5 (a) and 20 (b) occurrence records. Cells not meeting these criteria are not shown.

## References

Abrantes YG, Medeiros LS, Bennemann ABA et al (2020) Geographic distribution and conservation of seasonal killifishes (Cyprinodontiformes, Rivulidae) from the mid-northeastern caatinga ecoregion, northeastern Brazil. Neotrop Biol Conserv 15:301–315. 10.3897/neotropical.15.e51738

Abrantes YG, Ramos, TPA, Bento DDM, Lima SMQ (2023) Molecular delimitation of the seasonal killifishes of the *Hypsolebias antenori* species group (Cyprinodontiformes, Rivulidae), with description of two new species from the Caatinga coastal basins, northeastern Brazil. Zootaxa 5389:545–562. 10.11646/zootaxa.5389.5.2

Abrantes YG, Berbel-Filho WM, Carvalho RA, Ramos TPA, Lima SMQ (2026) Description of two threatened and polymorphic seasonal killifish species of the genus *Hypsolebias* (Cyprinodontiformes: Rivulidae) from the Piranhas-Açu River basin, in the Brazilian semiarid. Neotrop Ichthyol 24:e250044. 10.1590/1982-0224-2025-0044

Alonso F, Terán GE, Serra Alanís WS et al (2023) From the mud to the tree: phylogeny of *Austrolebias* killifishes, new generic structure and description of a new species (Cyprinodontiformes: Rivulidae). Zool J Linn Soc 199:280–309. 10.1093/zoolinnean/zlad032

Alonso F, Terán GE, Calviño P et al (2024) Expect the unexpected: a new species of killifish from a highly stochastic temporary wetland near Iguazú Falls (Cyprinodontiformes: Rivulidae). Can J Zool 102:298–314. 10.1139/cjz-2023-013

Alonso F, Terán GE, Serra Alanís WS et al (2025) The rise of a Titan: a new species of the giant *Titanolebias* killifishes, and its phylogeny (Cyprinodontiformes: Rivulidae). Zool Anz 316:253–265. 10.1016/j.jcz.2025.04.009

Arezo MJ, Papa NG, Berois N, Clivio G, Montagne J, De la Piedra S (2017) Annual killifish adaptations to ephemeral environments: Diapause i in two *Austrolebias* species. Dev Dyn 246:848–857. 10.1002/dvdy.24580

Austin KG, Elsen PR, Coronado ENH et al (2025) Mismatch Between Global Importance of Peatlands and the Extent of Their Protection. Conserv Lett 18:e13080. 10.1111/conl.13080

Berois N, Arezo MJ, Papa NG, Clivio GA (2012) Annual fish: developmental adaptations for an extreme environment. WIREs Dev Biol 1:595–602. 10.1002/wdev.39

Betts J, Young RP, Hilton-Taylor C et al (2020) A framework for evaluating the impact of the IUCN Red List of threatened species. Conserv Biol 34:632–643. 10.1111/cobi.13454

Boettiger C, Lang DT, Wainwright PC (2012) Rfishbase: Exploring, manipulating and visualizing FishBase data from R. J Fish Biol 81:2030–2039. 10.1111/j.1095-8649.2012.03464.x

Borgelt J, Dorber M, Høiberg MA et al. (2022) More than half of data deficient species predicted to be threatened by extinction. Commun Biol 5:679. 10.1038/s42003-022-03638-9

Borges AKM, Oliveira TPR, Rosa IL et al (2021) Caught in the (inter) net: online trade of ornamental fish in Brazil. Biol Conserv 263:109344. 10.1016/j.biocon.2021.109344

Borges AKM, Oliveira TPR, Alves RRN (2022) Marine or freshwater the role of ornamental fish keeper’s preferences in the conservation of aquatic organisms in Brazil. PeerJ 10:1–23. 10.7717/peerj.14387

Brooks TM, Bakarr MI, Boucher T et al (2004) Coverage provided by the global protected-area system: Is it enough? BioScience 12:1081–1091. 10.1641/0006-3568(2004)054[1081:CPBTGP]2.0.CO;2

Bürkner PC (2017) brms: An R package for Bayesian multilevel models using Stan. J Stat Softw 80. 10.18637/jss.v080.i01

Calhoun AJK, Mushet DM, Bell KP, Boix D, Fitzsimons JA, Isselin-Nondedeu F (2017) Temporary wetlands: challenges and solutions to conserving a ‘disappearing’ ecosystem. Biol Conserv 211:3–11. 10.1016/j.biocon.2016.11.024.

Carvalho RL, Resende AF, Barlow J et al (2023) Pervasive gaps in Amazonian ecological research. Curr Biol 33:3495–3504. 10.1016/j.cub.2023.06.077

Castro RMC, Polaz CNM (2020) Small-sized fish: the largest and most threatened portion of the megadiverse Neotropical freshwater fish fauna. Biota Neotrop 20:e20180683. 10.1590/1676-0611-bn-2018-0683

Cazalis V, Santini L, Lucas PM et al (2023) Prioritizing the reassessment of data-deficient species on the IUCN Red List. Conserv Biol 37:e14139. 10.1111/cobi.14139

Clouser PR, Riggs CL, Romney ALT, Podrabsky JE (2025) Diapause and Anoxia-Induced Quiescence Are Unique States in Embryos of the Annual Killifish, *Austrofundulus limnaeus*. Biomolecules 15:515. 10.3390/biom15040515

Costa JHA, Zuanon J, Selinger A, Castilho AG, Souza UP, Duarte RM (2025) Diving into temporary pools: a global perspective on freshwater fish research. Neotrop Ichthyol 23:e250043. 10.1590/1982-0224-2025-0043

Costa WJEM (2009) Aplocheiloid fishes of the Brazilian Atlantic Forest: history, diversity and conservation. Museu Nacional, Rio de Janeiro

Costa WJEM (2011). Identity of *Rivulus ocellatus* and a new name for a hermaphroditic species of *Kryptolebias* from south-eastern Brazil (Cyprinodontiformes: Rivulidae). Ichthyological Exploration of Freshwaters, 22:185.

Costa WJEM (2012) Delimiting priorities while biodiversity is lost: Rio’s seasonal killifishes on the edge of survival. Biodivers Conserv 21:2443–2452. 10.1007/s10531-012-0301-7

Costa WJEM (2019) Description of a new species of cynopoeciline killifish (Cyprinodontiformes, Aplocheilidae), possibly extinct, from the Atlantic Forest of south-eastern Brazil. ZooKeys 867:73–85. 10.3897/zookeys.867.34034

Costello MJ, May RM, Stork NE (2013) Can we name earth’s species before they go extinct? Science 339:413–416. 10.1126/science.1230318

Deguignet M, Arnell A, Juffe-Bignoli D et al (2017) Measuring the extent of overlaps in protected area designations. PLoS One 12:e0188681. 10.1371/journal.pone.0188681

Diniz-Filho JAF, Jardim L, Guedes JJM et al (2023) Macroecological links between the Linnean, Wallacean, and Darwinian shortfalls. Front Biogeogr 15:e59566. 10.21425/F5FBG59566

Drawert HA (2022) A new species of the seasonal killifish genus *Moema* (Cyprinodontiformes: Rivulidae) from the Piraí watershed in the Southwest Amazon basin. Neotrop Ichthyol 20: e220067. 10.1590/1982-0224-2022-0067

Drawert HA (2023) Rivulids (Rivulidae: Cyprinodontiformes) of Bolivia: knowledge status and updated inventory. Neotrop Hydrobiol Aquat Conserv 4:1–83. 10.55565/nhac.zlmk9566

Drawert HA, Ergueta C (2024) Redescription of *Austrolebias accorsii* (Cyprinodontiformes: Rivulidae) and description of a new species of the genus from the upper Paraguay River basin. Neotrop Ichthyol 22:e240001. 10.1590/1982-0224-2024-0001

Dudley N (2008) Guidelines for Applying Protected Area Management Categories. IUCN, Gland, Switzerland.

Faurby S, Farooq H (2026) Global patterns in species description year cannot reliably estimate undescribed diversity. Front Biogeogr 19:e171197. 10.21425/fob.19.171197

Frateles LEF, da Silva NJ, Terribile LC, Diniz-Filho JAF (2024) Linnean shortfall and space-time patterns in species description of New World coralsnakes (Serpentes: Elapidae). Zool Scr 53:299–311. 10.1111/zsc.12644

Frateles L, Ronquillo C, Nakamura G, Diniz-Filho JAF, Ladle RJ, Hortal J (2026) Scrutinizing the Wallacean shortfall: global gaps in snake occurrence data across space and environment. Ecography e08589. 10.1002/ecog.08589

Freitas TMS, Stropp J, Calegari BB et al (2021) Quantifying shortfalls in the knowledge on Neotropical Auchenipteridae fishes. Fish Fish 22:87–104. 10.1111/faf.12507

Fricke R, Eschmeyer WN, Van der Laan R (2026a) ESCHMEYER’S CATALOG OF FISHES: GENERA, SPECIES, REFERENCES. http://researcharchive.calacademy.org/research/ichthyology/catalog/fishcatmain.asp. Accessed June 2026.

Fricke R, Eschmeyer WN, Fong JD (2026b) Species by family/subfamily. http://researcharchive.calacademy.org/research/ichthyology/catalog/SpeciesByFamily.asp. Accessed 03 June 2026.

Froese R, Pauly D (2026) Fishbase. World Wide Web electronic publication. www.fishbase.org. Accessed 03 June 2026.

Furness AI (2016). The evolution of an annual life cycle in killifish: adaptation to ephemeral aquatic environments through embryonic diapause. Biol Rev 91:796–812. 10.1111/brv.12194

Furness AI, Reznick DN, Tatarenkov A, Avise JC (2018) The Evolution of Diapause in *Rivulus* (*Laimosemion*). Zool J Linn Soc 18:773–790. 10.1093/zoolinnean/zly021

GBIF.org (2026) GBIF Occurrence Download. 10.15468/dl.as44ck

Gearty W, Chamberlain S (2025) rredlist: ‘IUCN’ Red List Client. R package version 1.1.1. 10.32614/CRAN.package.rredlist

Geldmann J, Manica A, Burgess ND, Coad L, Balmford A (2019) A global-level assessment of the effectiveness of protected areas at resisting anthropogenic pressures. PNAS 116:23209–23215. 10.1073/pnas.1908221116

Godoy RS, Weber V, Lanés LEK et al (2025a) Early post-hatching sensitivity of a Neotropical annual killifish to glyphosate-based herbicide. Toxicol Environ Chem 107:1322–1338. 10.1080/02772248.2025.2527648.

Godoy RS, Weber V, Hoffmann P et al (2025b) Topsoil translocation as a restoration tool for endangered seasonal killifish in temporary wetlands. Restor Ecol 34:e70262. 10.1111/rec.70262

Gombin J, Vaidyanathan R, Agafonkin V (2025) concaveman: A Very Fast 2D Concave Hull Algorithm. R package version 1.2.0. https://github.com/joelgombin/concaveman

Guedes GHS, Gomes ID, do Nascimento AA et al (2023a) Reproductive strategy of the annual fish *Leptopanchax opalescens* (Rivulidae) and trade-off between egg size and maximum body length in temporary wetlands. Wetlands, 43:29. 10.1007/s13157-023-01680-9

Guedes GHS, Luz CHP, Mazzoni R, Lira FO, Araújo FG (2023b) New occurrences of the endangered *Notholebias minimus* (Cyprinodontiformes: Rivulidae) in coastal plains of the state of Rio de Janeiro, Brazil: populations features and conservation. Neotrop Ichthyol 21:e230013. 10.1590/1982-0224-2023-0013

Guedes GHS, Santangelo JM, Pires APF, Araújo FG (2025a) β-Diversity Scaling Patterns Across Different Bioregionalisations for a Megadiverse Neotropical Fish Family. J Biogeogr 52: e15088. 10.1111/jbi.15088

Guedes GHS, Luz CHP, Souza VJ, Araújo FG (2025b) A fish frontier? Itatiaia expedition and biodiversity repositories reveal gaps in fish occurrences in Brazil’s high-altitude aquatic ecosystems. Zoologia, 42:e24077. 10.1590/S1984-4689.v42.e24077

Guedes GHS, Araújo FG (2026) Water and fire: Wildfires threaten fish habitats across Brazilian biomes, and protected areas offer insufficient safeguards. Biol Conserv 315:111692. 10.1016/j.biocon.2025.111692

Guedes GHS, Cordeiro L, Azeredo, LFSP, Araújo FG (2026b) Microplastics in wetlands: contrasting fish contamination between mangroves and temporary ponds in southeastern Brazil. Wetl Ecol Manag 34:49. 10.1007/s11273-026-10156-6

Guedes JJM, Gomes De Lima HV, Mendonça LR, Chen-Zhao R, Diniz-Filho JAF, Moura MR (2024) Temporal trends in global reptile species descriptions over three decades. Syst Biodivers 22. 10.1080/14772000.2024.2419832

Guedes JJM, Santos JP, Freitas AVL (2026a) Unveiling Shortfalls in Biodiversity Knowledge of Brazilian Butterflies. Diversity 18:373. 10.3390/d18060373

Guerra V, Jardim L, Llusia D, Márquez R, Bastos RP (2020) Knowledge status and trends in description of amphibian species in Brazil. Ecol Indic 118: 106754. 10.1016/j.ecolind.2020.106754

Hollister JW (2025) elevatr: Access Elevation Data from Various APIs. R package version 0.99.1. https://CRAN.R-project.org/package=elevatr/

Hortal J, De Bello F, Diniz-Filho JAF, Lewinsohn TM, Lobo JM, Ladle RJ (2015) Seven Shortfalls that Beset Large-Scale Knowledge of Biodiversity. Annu Rev Ecol Syst 46:523–549. 10.1146/annurev-ecolsys-112414-054400

Hsieh TC, Ma KH, Chao A (2024) iNEXT: iNterpolation and EXTrapolation for species diversity. R package version 3.0.1. http://chao.stat.nthu.edu.tw/wordpress/software-download

ICMBio (2025a) Sumário Executivo – Plano de Ação Nacional para Conservação dos Peixes Rivulídeos Ameaçados de Extinção. Pirassununga. https://www.gov.br/icmbio/pt-br/assuntos/biodiversidade/pan/pan-rivulideos/2-ciclo/pan-rivulideos-sumario.pdf

ICMBio (2025b) Diretrizes para adequada consideração dos peixes rivulídeos nos empreendimentos passíveis de atos autorizativos ambientais. Plano de Ação Nacional para Conservação dos Peixes Rivulídeos Ameaçados de Extinção. Pirassununga. https://www.gov.br/icmbio/pt-br/assuntos/biodiversidade/pan/pan-rivulideos

ICMBio (2026a). Sistema de Avaliação do Risco de Extinção da Biodiversidade – SALVE. https://salve.icmbio.gov.br. Accessed 19 June 2026.

ICMBio (2026b) Avaliação de Manejo Ex situ de espécies do gênero *Steindachneridion* e *Ophthalmolebias constanciae*. ICMBio, Brasília.

IUCN (2026) The IUCN Red List of Threatened Species. Version 2025–2. https://www.iucnredlist.org

Jarvis A, Reuter HI, Nelson A, Guevara E (2008) Hole-filled SRTM for the globe Version 4. CGIAR-CSI SRTM 90m Database. https://srtm.csi.cgiar.org

Joppa LN, Pfaff A (2009) High and far: Biases in the location of protected areas. PLoS ONE, 4:e8273. 10.1371/journal.pone.0008273

Joppa LN, Roberts DL, Pimm SL (2011) How many species of flowering plants are there? Proc R Soc B: Biol Sci 278:554–559. 10.1098/rspb.2010.1004

Lehner B, Grill G (2013) Global river hydrography and network routing: Baseline data and new approaches to study the world’s large river systems. Hydrol Process 27:2171–2186. 10.1002/hyp.9740

Li G, Fang C, Watson JEM et al (2024) Mixed effectiveness of global protected areas in resisting habitat loss. Nat Commun 15:8389. 10.1038/s41467-024-52693-9

Lima M, Freitas TMS, Brasil LS et al (2026) Unveiling the Global Linnean Shortfall in Leptophlebiidae (Ephemeroptera, Insecta): Historical Trends and Conservation Implications. Aquat Conserv: Mar Freshw Ecosyst 36:e70402. 10.1002/aqc.70402

Loureiro M, de Sá R, Serra SW et al (2018). Review of the family Rivulidae (Cyprinodontiformes, Aplocheiloidei) and a molecular and morphological phylogeny of the annual fish genus *Austrolebias* Costa 1998. Neotrop Ichthyol 16:e180007. 10.1590/1982-0224-20180007

Lu M, He F (2017) Estimating regional species richness: The case of China’s vascular plant species. Glob Ecol Biogeogr 26:835–845. 10.1111/geb.12589

Mace GM (2004) The role of taxonomy in species conservation. Philos Trans R Soc B Biol Sci 359:711–719. 10.1098/rstb.2003.1454

Mammola S, Adamo M, Antić D et al (2023) Drivers of species knowledge across the tree of life. Elife 12. 10.7554/elife.88251

Massicotte P, South A (2023) rnaturalearth: World Map Data from Natural Earth. R package version 1.0.1. https://CRAN.R-project.org/package=rnaturalearth

Meyer L, Ladle R, Trad R et al (2026) Taxonomic uncertainty: causes, consequences, and metrics. Trends Ecol Evol 41:299–308. 10.1016/j.tree.2025.12.006

Miqueleiz I, Bohm M, Ariño AH, Miranda R (2020) Assessment gaps and biases in knowledge of conservation status of fishes. Aquatic Conserv: Mar Freshw Ecosyst 30:225–236. 10.1002/aqc.3282

Mora C, Tittensor DP, Adl S, Simpson AGB, Worm B (2011) How many species are there on Earth and in the ocean? PLoS Biol 9:e1001127. 10.1371/journal.pbio.1001127

Mouquet N, Langlois J, Casajus N et al (2024) Low human interest for the most at-risk reef fishes worldwide. Sci Adv 10:eadj9510. 10.1126/sciadv.adj9510

Moura MR, Costa HC, Peixoto MA, Carvalho ALG, Santana DJ, Vasconcelos HL (2018) Geographical and socioeconomic determinants of species discovery trends in a biodiversity hotspot. Biol Conserv 220:237–244. 10.1016/j.biocon.2018.01.024

Moura MR, Jetz W (2021). Shortfalls and opportunities in terrestrial vertebrate species discovery. Nat Ecol Evol 5:631–639.

Nakamura G, Stabile BHM, Frateles LEF et al (2025) The hidden biodiversity knowledge split in biological collections. Proc R Soc B: Biol Sci 292:20251045. 10.1098/rspb.2025.1045

Nielsen DTB, Hoetmer JW, Vandekerckhove E (2025) *Moema humaita*, a new species of annual fish (Cyprinodontiformes: Rivulidae) from the middle rio Madeira, Amazon basin, Brazil. Zootaxa 5631:189–197. 10.11646/zootaxa.5631.1.10

Oliveira U, Paglia AP, Brescovit AD et al (2016) The strong influence of collection bias on biodiversity knowledge shortfalls of Brazilian terrestrial biodiversity. Divers Distrib 22:1232–1244. 10.1111/ddi.12489

Pavanelli CS, Fukakusa CK, Da Costa Neto FPS et al (2024a) Kryptolebias ocellatus. ICMBio. 10.37002/salve.ficha.24966.2

Pavanelli CS, Fukakusa CK, Da Costa Neto FPS et al (2024b) Simpsonichthys zonatus. ICMBio. 10.37002/salve.ficha.25026.2

Pearson DL, Hamilton AL, Erwin TL (2011) Recovery plan for the endangered taxonomy profession. BioScience 61:58–63. 10.1525/bio.2011.61.1.11

Pebesma E (2018) Simple features for R: Standardized support for spatial vector data. R J 10:439–446. 10.32614/rj-2018-009

Penhacek M, Castro-Souza RA, Tessarolo G, Diniz-Filho JA, Sobral-Souza T, de Jesus Rodrigues D (2025) Biases in Amphibian Sampling in the Amazon: Using Infrastructure and Accessibility Data to Identify Sampling Gaps. Biotropica 57:e70079. 10.1111/btp.70079

Pinheiro J, Bates D, DebRoy S, Sarkar D, Heisterkamp S, Van Willigen B (2020) Package nlme: Linear and Nonlinear Mixed Effects Models. R Package, 1–336. https://cran.r-project.org/web/packages/nlme/nlme.pdf

QGIS Development Team (2026) QGIS Geographic Information System. Chicago, IL: Open Source Geospatial Foundation Project.

Ramos TPA, Nielsen DTB, Abrantes YG, Lira FO, Lustosa-Costa SY (2023) A new species of cloud fish of the genus *Hypsolebias* from Northeast Brazil (Cyprinodontiformes: Rivulidae). Neotrop Ichthyol 21:e230068. 10.1590/1982-0224-2023-0068

Reis GS, Tejerina-Garro FL, Dagosta FCP, Teresa FB, Carvalho RA (2024) Seeking for gaps in taxonomic descriptions of endemic fishes: a pathway to challenge the Linnean shortfall in a Neotropical basin. Neotrop Ichthyol 22:e230128. 10.1590/1982-0224-2023-0128

Rodrigues ASL, Andelman SJ, Bakan MI et al (2004) Effectiveness of the global protected area network in representing species diversity. Nature, 428:640–643. 10.1038/nature02422

Sarmento-Soares LM, Vieira-Guimarães F, Martins-Pinheiro RF (2026) Patterns of Diversity and Endemism of Killifishes (Cyprinodontiformes: Rivulidae) in the Southeastern and Eastern Coastal Basins of the Atlantic Forest, Brazil. Diversity 18:317. 10.3390/d18060317

Sayer CA, Fernando E, Jimenez RR et al (2025) One-quarter of freshwater fauna threatened with extinction. Nature 638:138–145. 10.1038/s41586-024-08375-z

Souza ECF, Brant A, Rangel CA et al (2018) Avaliação do risco de extinção da fauna brasileira: ponto de partida para a conservação da biodiversidade. Divers Gest 2:62–75.

Tapley B, Michaels CJ, Gumbs R et al (2018) The disparity between species description and conservation assessment: a case study in taxa with high rates of species discovery. Biol Conserv 220:209–214. 10.1016/j.biocon.2018.01.022

Tedesco PA, Bigorne R, Bogan AE, Giam X, Jézéquel C, Hugueny B (2014) Estimating how many undescribed species have gone extinct. Conserv Biol 28:1360–1370. 10.1111%2Fcobi.12285

UNEP-WCMC, IUCN (2026) Protected Planet: The World Database on Protected Areas (WDPA) and World Database on Other Effective Area-based Conservation Measures (WD-OECM) [Online]. Cambridge, UK: UNEP-WCMC and IUCN. www.protectedplanet.net

Volcan MV, Lanés LEK (2018) Brazilian killifishes risk extinction. Science 361:340341. 10.1126/science.aau5930

Volcan MV, Severo-Neto F, Lanés LEK (2018) Unrecognized biodiversity in a world’s hotspot: Three new species of *Melanorivulus* (Cyprinodontiformes: Rivulidae) from tributaries of the right bank of the Rio Paraná basin, Brazilian Cerrado. Zoosyst Evol 94:263–280. 10.3897/zse.94.24406

Volcan MV, Suárez YR, Severo-Neto F, Amorim PF, Costa WJEM (2024) A typical coastal plain taxon in central Brazil: relationships and description of a new species of non-annual killifish (Cyprinodontiformes; Rivulidae) from the Paraná River basin. Zool Anz 312:92–102. 10.1016/j.jcz.2024.07.013

Volcan MV, Garcez DK, Robe LJ, Feltrin CRM, Costa WJEM, Lanés LEK (2025) A new and threatened species of internally inseminating seasonal killifish of *Campellolebias* (Cyprinodontiformes: Rivulidae) endemic to a continental island in the Atlantic Forest, Southern Brazil. Zool Anz 316:75–84. 10.1016/j.jcz.2025.03.00

Zapata FA, Robertson DR (2007) How many species of shore fishes are there in the Tropical Eastern Pacific? J Biogeogr 34:38–51. 10.1111/j.1365-2699.2006.01586.x

Zajic DE, Podrabsky JE (2020) Metabolomics analysis of annual killifish (*Austrofundulus limnaeus*) embryos during aerial dehydration stress. Physiol Genomics 52:408–422. 10.1152/physiolgenomics.00072.2020

Zizka A, Silvestro D, Andermann T et al (2019) CoordinateCleaner: Standardized cleaning of occurrence records from biological collection databases. Methods Ecol Evol 10:744–751. 10.1111/2041-210X.13152

